# Microglial CD68 and L-ferritin upregulation in response to phosphorylated-TDP-43 pathology in the amyotrophic lateral sclerosis brain

**DOI:** 10.1101/2023.02.14.528561

**Authors:** Molly E V Swanson, Miran Mrkela, Helen C Murray, Maize C Cao, Clinton Turner, Maurice A Curtis, Richard L M Faull, Adam K Walker, Emma L Scotter

## Abstract

Microglia, the innate immune cells of the brain, are activated by damage or disease. In mouse models of amyotrophic lateral sclerosis (ALS), microglia shift from neurotrophic to neurotoxic states with disease progression. It remains unclear how human microglia change relative to the TAR DNA-binding protein 43 (TDP-43) aggregation that occurs in 97% of ALS cases. Here we examine spatial relationships between microglial activation and TDP-43 pathology in brain tissue from people with ALS and from a TDP-43-driven ALS mouse model. Post-mortem human brain tissue from the Neurological Foundation Human Brain Bank was obtained from 10 control and 10 ALS cases in parallel with brain tissue from a bigenic *NFFH*-tTA/*tetO*-hTDP-43ΔNLS (rNLS) mouse model of ALS at disease onset, early disease, and late disease stages. The spatiotemporal relationship between microglial activation and ALS pathology was determined by investigating microglial functional marker expression in brain regions with low and high TDP-43 burden at end-stage human disease: hippocampus and motor cortex, respectively. Sections were immunohistochemically labelled with a two-round multiplexed antibody panel against; microglial functional markers (L-ferritin, HLA-DR, CD74, CD68, and Iba1), a neuronal marker (NeuN), an astrocyte marker (GFAP), and pathological phosphorylated TDP-43 (pTDP-43). Single-cell levels of microglial functional markers were quantified using custom analysis pipelines and mapped to anatomical regions and ALS pathology. We identified a significant increase in microglial Iba1 and CD68 expression in the human ALS motor cortex, with microglial CD68 being significantly correlated with pTDP-43 pathology load. We also identified two subpopulations of microglia enriched in the ALS motor cortex that were defined by high L-ferritin expression. A similar pattern of microglial changes was observed in the rNLS mouse, with an increase first in CD68 and then in L-ferritin expression, with both occurring only after pTDP-43 inclusions were detectable. Our data strongly suggest that microglia are phagocytic at early-stage ALS but transition to a dysfunctional state at end-stage disease, and that these functional states are driven by pTDP-43 aggregation. Overall, these findings enhance our understanding of microglial phenotypes and function in ALS.

## Introduction

Amyotrophic lateral sclerosis (ALS) is the most common form of motor neuron disease. ALS is characterised by progressive death of upper and lower motor neurons, which leads to muscle atrophy, disability, and death [1]. Diverse genetic and environmental factors cause or increase the risk of ALS [2,3]. Despite this range of risk factors, 97% of all ALS cases share a common pathological hallmark: intracellular accumulation of TAR DNA-binding protein 43 (TDP-43) aggregates in motor neurons [4]. Motor neurons are central to ALS pathogenesis and symptomatology but microglia, the innate immune cells of the brain, are also altered in disease. Microglia may become reactive in response to damage or disease pathology to maintain brain homeostasis [5]. However, evidence from rodent *SOD1* models of ALS indicates that some microglial functional states could be harmful later in disease and contribute to ALS progression [6]. It remains unclear how microglia change spatiotemporally relative to TDP-43 pathology in ALS and how microglial function impacts neurodegeneration.

Microglial functions are altered in a range of neurodegenerative diseases, with changes in microglial morphology and protein expression identified in proximity to pathological protein aggregates [7]. These changes in microglial function are hypothesised to occur to protect the brain from degeneration, but also have the potential to contribute to neurodegeneration. In the G93A *SOD1* ALS mouse model, isolated primary microglia at an early symptomatic stage were neuroprotective, supporting the growth of motor neurons in co-culture [8]. Conversely, primary microglia isolated from the same model at a late symptomatic stage were neurotoxic, inducing motor neuron death [8]. Therefore, microglia are considered to have a biphasic role in *SOD1*-linked ALS, supporting motor neuron survival early in disease but later switching to promote disease pathogenesis. The phenotype of microglia in ALS with TDP-43 pathology is less well characterised. A neurotoxic microglial phenotype can be induced by native and mutant forms of TDP-43 *in vitro* [9], demonstrating the capacity for a microglial functional shift in response to TDP-43. Furthermore, specific microglial states present in the human ALS spinal cord containing pTDP-43 pathology negatively correlated with disease progression [10]. In contrast, the nuclear-localisation sequence-defective TDP-43 mouse model of ALS demonstrated microglia express unique suites of genes at each disease state, with strong neuroprotective gene signatures being identified at early- and late-stage disease, and recovery, arguing against a late-stage neurotoxic microglial phenotype [11].

Underpinning the neuroprotective, neurotoxic, and other functional phenotypes of microglia are changes in protein expression of key functional proteins. L-ferritin, HLA-DR, CD68, CD74, and Iba1 are microglial proteins that report upon a range of microglial functions and are altered across various human ALS genotypes and models of ALS (Table S1) [12–28]. L-ferritin is a marker of microglial dysfunction [29,30], HLA-DR and CD74 are involved in antigen presentation and are classical markers of ‘activation’ [31–35], CD68 is expressed in lysosomes and reports on phagocytosis [36,37], and Iba1 labels the cytoskeleton and can report on morphological changes [38–40]. Microglial Iba1 and CD68 expression are increased in human ALS brain regions with high TDP-43 pathological load and neuronal loss [41]. Furthermore, CD68 expression in ALS is strongly correlated with motor cortex phosphorylated TDP-43 (pTDP-43) pathology load [25,42], disease progression, and upper motor neuron clinical symptoms [43,44]. However, several of these human studies lacked age-matched control cases [41,43] and the small number of microglial markers examined leave unclear whether microglial functional changes in response to TDP-43 pathology are neuroprotective or neurotoxic.

As evidenced above, microglial functional states in post-mortem tissues have typically been inferred from cell morphology and/or protein expression of individual microglial proteins or limited sets of proteins by single-label chromogenic immunohistochemistry. While traditional immunohistochemical studies provide proteomic data with spatial context, single chromogen or even triple fluorescent labelling methods do not capture microglial heterogeneity [30]. Indeed, microglial research is moving away from rigid categorisation of a limited number of distinct functional states, and towards a multidimensional integration of transcriptomic, proteomic, metabolomic, epigenetic, and morphological data to define microglial phenotypes [45]. While bulk tissue RNA sequencing and single cell-based methodologies like single cell- and single nuclear-RNA sequencing have provided high resolution transcriptomic information on microglial changes in ALS, these datasets lack proteomic information and spatial context [18,20,21,26–28,46–49]. Further, although the identification of microglial changes in post-mortem human ALS brain tissue is key to identify disease-relevant therapeutic targets, this tissue is collected at end-stage disease. Current methods of analysing microglial changes in human ALS tissue thus fail to account for complex protein expression changes and preclude the assessment of such changes over time, including in early stages of disease progression that are critical for understanding disease mechanisms.

To address these shortcomings, here we utilise multiplexed immunohistochemistry and single-cell analysis to compare the protein expression of HLA-DR, CD68, CD74, and Iba1 between ALS brains and age-matched normal controls. We also analyse the spatial relationship between individual microglial phenotypes and pTDP-43 pathology in the ALS brain. And finally, we infer temporal changes in human ALS by examining human brain regions affected with neuropathology both early (motor cortex) and late (hippocampus) disease stages [50], and TDP-43 mouse model brain regions at disease onset, early disease, and late disease stages [51]. By analysing post-mortem tissue from patients with end-stage ALS, in parallel with tissue from rNLS mice at different disease time points, we provide an assessment of microglial changes relative to pTDP-43 in the human brain with temporal context.

## Methods

### Human tissue selection

Formalin-fixed paraffin-embedded motor cortex and hippocampus blocks from 10 neurologically normal and 10 ALS cases from the Neurological Foundation Human Brain Bank (HuBB) at the Centre for Brain Research, University of Auckland were used in this study (Table 1). Control cases had no previous history of neurological disorders and cause of death was unrelated to any neurological condition. ALS cases were diagnosed clinically by consultant neurologists at Auckland City or Middlemore Hospitals (Auckland, New Zealand) during life. All case classifications were confirmed with post-mortem neuropathology assessments performed by consultant neuropathologists at Auckland City Hospital. Control cases showed no disease pathology other than normal age-related amyloid deposition. All ALS cases showed pTDP-43 deposition in the motor cortex.

**Table 1.** Post-mortem tissue donor demographics.

| Table 1 Post-mortem tissue donor demographics |  |  |  |  |  |  |
| --- | --- | --- | --- | --- | --- | --- |
| Group |  | Case code | Age (y) | Sex | PMD (h) | fALS/sALS |
| Neurologically Normal Controls<br>(n = 10) |  | H211 | 41 | M | 8 | - |
|  |  | H215 | 67 | F | 23.5 | - |
|  |  | H226 | 73 | F | 48 | - |
|  |  | H230 | 57 | F | 32 | - |
|  |  | H238 | 63 | F | 16 | - |
|  |  | H240 | 73 | M | 26.5 | - |
|  |  | H242 | 61 | M | 19.5 | - |
|  |  | H245 | 63 | M | 20 | - |
|  |  | H247 | 51 | M | 31 | - |
|  |  | H249 | 77 | F | 17 | - |
| Control mean ± SD |  | - | 62.60 ± 10.95 | 5M:5F | 24.15 ± 11.06 | - |
| ALS<br>(n = 10) | Stage 1-3<br>(n = 5) | MN11 | 77 | F | 18 | fALS<br>(unknown) |
|  |  | MN14 | 59 | F | 19 | fALS<br>(unknown) |
|  |  | MN16 | 69 | M | 16.5 | sALS |
|  |  | MN20 | 85 | M | 15 | sALS |
|  |  | MN25 | 71 | M | 23.5 | sALS |
|  | Stage 4<br>(n = 5) | MN13 | 55 | M | 10 | sALS |
|  |  | MN15 | 54 | F | 18 | sALS |
|  |  | MN19 | 75 | M | 20.5 | sALS |
|  |  | MN29 | 72 | M | 24 | sALS |
|  |  | MN30 | 84 | F | 17.5 | sALS |
| ALS mean ± SD |  | - | 70.10 ± 11.07 | 6M:4F | 18.20 ± 4.063 | - |

Abbreviations: PMD, Post-mortem delay; fALS, familial ALS; sALS, sporadic ALS; SD, standard deviation; M, male; F, female; ALS, amyotrophic lateral sclerosis.

The fixation and dissection of anatomical regions from human brains by the HuBB has been described previously [52]. Briefly, human brains were obtained at autopsy and the right hemisphere was fixed by perfusion of 15% formaldehyde in 0.1 M phosphate buffer through the cerebral arteries. Brains were subsequently dissected into approximately 60 regional blocks and 1-cm thick blocks from each region were processed for paraffin embedding. For this study, 10-μm thick sections were cut from paraffin-embedded motor cortex and hippocampal blocks from all control and ALS cases (Table 1) using a rotary microtome and mounted on ÜberFrost^®^ Printer Slides (InstrumeC).

### Mouse tissue collection

rNLS double-transgenic experimental mice and monogenic litter-mate controls were produced from crosses of hemizygous B6.C3-Tg(NEFH-tTA)8Vle/JAusb mice (*NEFH-tTA* line 8, stock #025397) with homozygous B6.C3-Tg(*tetO*-TARDBP*)4Vle/JAusb mice (*tetO*-hTDP-43^ΔNLS^ line 4, stock #014650), originally obtained from Jackson Laboratories [51]following backcrossing of each independent line for >10 generations on C56BL/6JAusb (Australian BioResources) background. Intercross mice and offspring were provided with 200mg/kg doxycycline (Dox)-containing feed (Specialty Feeds, Australia) until 8-12 weeks of age. Both sexes were included in each group, and mice were housed under 12-h light/dark cycle at 21 ± 1 °C and 55 ± 5% humidity. Brains were collected from mice as described previously [53]. Briefly, at pre-determined timepoints after removal of Dox-containing feed, mice were deeply anaesthetized and transcardially perfused with phosphate-buffered saline followed by 4% paraformaldehyde. Brains were post-fixed in 4% PFA for ~16 hours, dehydrated through increasing concentrations of ethanol and embedded in paraffin. Tissues sections (10 μm thickness) were mounted on StarFrost slides. Animal ethics approval was obtained from The University of Queensland (#QBI/131/18), and experiments were conducted in accordance with the Australian code of practice for the care and use of animals for scientific purposes.

### Paraffin immunohistochemistry

Immunohistochemical labelling panels were designed to investigate changes in microglial functional protein expression relative to ALS neurodegeneration in the human and mouse brains. The panels consisted of 2 rounds of immunohistochemical labelling encompassing microglial functional markers and anatomical and neuropathology markers (Table 2).

**Table 2.** Multiplex immunohistochemistry panels for analysis of microglial changes in the human ALS and hTDP-43-ΔNLS ‘rNLS’ mouse brains.

| PANEL 1 Human |  |  |  |
| --- | --- | --- | --- |
| Round 1 primary antibody | Round 1 secondary antibody | Round 2 primary antibody | Round 2 secondary antibody |
| Mouse IgG<sub>2b</sub> anti-L-ferritin<br>(1:1,000; Abcam, Ab218400) | Goat anti-mouse IgG<sub>2b</sub> AlexaFluor® 488<br>(1:500; ThermoFisher, A21141) | Mouse IgG<sub>1</sub> anti-TDP-43, phospho Ser409/410<br>(1:1,000; Cosmobio, CAC-TIP-PTD-M01) | Goat anti-mouse IgG1 AlexaFluor® 488<br>(1:500; ThermoFisher, A21121) |
| Mouse IgG<sub>1</sub> anti-HLA-DQ, DP, DR<br>(1:500; DAKO, M0775) | Goat anti-mouse IgG<sub>1</sub> AlexaFluor® 546<br>(1:500; ThermoFisher, A21123) | Guinea pig anti-NeuN<br>(1:500; Sigma Aldrich, ABN90) | Goat anti-guinea pig AlexaFluor® 546<br>(1:500; ThermoFisher, A11074) |
| Mouse IgG<sub>3</sub> anti-CD68<br>(1:200; Abcam, Ab783) | Goat anti-mouse IgG<sub>3</sub> AlexaFluor® 594<br>(1:500; ThermoFisher, A21155) | Rabbit anti-GFAP<br>(1:2,000; DAKO, Z0334) | Goat anti-rabbit AlexaFluor® 594<br>(1:500; Jackson Laboratories, 111-586-144) |
| Rabbit anti-CD74<br>(1:1,000; Abcam, Ab64772) | Goat anti-rabbit AlexaFluor® 647<br>(1:500; Jackson Laboratories, 111-605-144) | UEA-1 Lectin, biotinylate<br>(1:1,000; Sigma-Aldrich, L8262) | Streptavidin AlexaFluor® 647<br>(1:500; Jackson Laboratories, 016-170-084) |
| Chicken anti-Iba1<br>(1:1,000; Synaptic Systems, 234009) | Goat anti-chicken AlexaFluor® 800-plus<br>(1:500; ThermoFisher, A32935) |  |  |
| PANEL 2 Mouse |  |  |  |
| Round 1 primary antibody | Round 1 secondary antibody | Round 2 primary antibody | Round 2 secondary antibody |
| Rabbit anti-CD68<br>(1:1,000; Abcam, Ab215212) | Donkey anti-rabbit IgG AlexaFluor® 488<br>(1:500; Jackson Laboratories, 711-545-152) | Rabbit anti-pTDP-43, Ser 409/410<br>(1:3,000; Cosmo Bio, TIP-PTD-P02) | Donkey anti-rabbit AlexaFluor® 488<br>(1:500; Jackson Laboratories, 711-545-152) |
| Goat anti-L-ferritin<br>(1:1,000; Lifespan Biosciences, LS-B4383-50) | Donkey anti-goat AlexaFluor® 546<br>(1:500; ThermoFisher, A11056) | - | - |
| Guinea pig anti-NeuN<br>(1:500; Sigma Aldrich, ABN90) | Donkey anti-guinea pig AlexaFluor® 594<br>(1:500; Jackson Laboratories, 706-585-148) | - | - |
| Chicken anti-Iba1<br>(1:1,000; Synaptic Systems, 234009) | Donkey anti-chicken AlexaFluor® 647<br>(1:500; Jackson Laboratories, 703-605-155) | Goat anti-Iba1<br>(1:1,000; Abcam, Ab5076) | Donkey anti-goat AlexaFluor® 647<br>(1:500; Jackson Laboratories, 705-605-947) |

Immunohistochemical labelling of paraffin-embedded motor cortex and hippocampal sections was carried out as previously described [54,55]. In short, Tris-EDTA pH 9.0 antigen retrieval was performed, and sections were incubated in primary then secondary antibody mixtures (Table 2). Nuclei were counterstained with Hoechst 33258 and sections were coverslipped using ProLong^®^ Gold Antifade mounting media.

After the first round of labelling, sections were imaged on a Zeiss Z2 Axioimager (10×/0.45 NA) using MetaSystems VSlide acquisition software and MetaCyte stitching software. The Zeiss Z2 Axioimager used a Colibri 7 solid-state light source with LED lamps and the following filter sets to enable spectral separation of the 6 fluorophores per round (Ex peak (nm); Em (nm)/bandpass (nm)): Hoechst 33258 (385; 447/60), AlexaFluor^®^ 488 (475; 550/32), AlexaFluor^®^ 546 (555; 580/23), AlexaFluor^®^ 594 (590; 628/32), AlexaFluor^®^ 647 (630; 676/29), and AlexaFluor^®^ 800 (735; 809/81).

Sections were subsequently decoverslipped and incubated in 5x LI-COR NewBlot IR Stripping Buffer for 13 minutes at room temperature to strip the round 1 primary and secondary antibodies [56,57]. Following stripping, sections were incubated in the round 2 primary then secondary antibody mixtures, counterstained with Hoechst, and coverslipped. The round 2 labelling was then imaged within the same area of tissue as for round 1.

Individual channel images were extracted, using VSViewer v2.1.112, from at least two areas of grey matter per motor cortex section and the entire dentate gyrus per hippocampal section. Images from round 1 and 2 labelling were aligned for analysis using the Hoechst 33258 nuclear counterstain, as previously described [54,57].

Absence of bleed-through between adjacent channels was confirmed by labelling for all primary antibodies in a given round (Table 2) and either Hoechst 33258 alone, Hoechst 33258 alone and all secondaries, or Hoechst 33258 and secondaries in alternating channels (488, 594, and 800 nm; or 546 and 647 nm) (Fig. S1). Absence of cross-reactivity between antibodies within a single round was also confirmed (Fig. S1). There was minor cross-reactivity between the mouse IgG1 anti-pTDP-43 from round 1 and the mouse IgG1 anti-HLA-DR in round 2 in some human cases, which was accounted for in subsequent analyses.

### Quantification using MetaMorph custom image analysis pipelines

To quantify microglial changes relative to neurodegeneration in the ALS brain, we developed custom image analysis pipelines in Metamorph software (Molecular Devices) similar to those previously described [57,58]. Three image analysis pipelines were developed for this study: tissue-wide pathology load analysis, and tissue-wide and single-cell microglial functional marker expression analysis (Fig. S2). Prior to all analyses, image processing was carried out; on a composite image of Hoechst and NeuN, manual regions of interest (ROI) were drawn to retain tissue areas for analysis and exclude tissue folds and defects. Motor cortex ROIs encompassed layers I-VI and hippocampus ROIs encompassed the molecular, granule cell, and polymorphic layers of the dentate gyrus.

#### Tissue-wide pathology load analyses

We first analysed the overall load of pTDP-43 and GFAP in control versus ALS tissues. To eliminate high HLA-DR immunoreactivity from the previous immunohistochemistry round in the human sections, a binary mask of high HLA-DR immunoreactivity was generated using the threshold clip tool and subtracted from the pTDP-43 image. To isolate specific pTDP-43 or GFAP immunoreactivity, binary masks of high immunoreactivity were generated for the HLA-DR-subtracted pTDP-43 image (or raw pTDP-43 in the mouse tissue) and the raw GFAP image using the threshold clip tool. The integrated intensity of the pTDP-43 labelling within the binary mask and area of the GFAP binary mask were then measured and normalised to the ROI area.

#### Tissue-wide microglial functional marker expression analysis

To identify global microglial changes in the ALS brain, we first measured the tissue-wide abundance of each microglial marker. In the human tissue, binary masks of Iba1, HLA-DR, CD68, and CD74 immunolabelling were generated using the adaptive thresholding tool. A binary mask of high L-ferritin immunoreactivity was not generated, as L-ferritin immunoreactivity was identified on microglia as well as GFAP-positive astrocytes (Fig. S3). The binary masks from all thresholded markers of interest were combined to create a master mask, encompassing all microglia immunoreactive for Iba1, HLA-DR, CD68, and/or CD74. In the mouse tissue, a binary mask of only Iba1 was generated to identify all microglia as all microglia were highly immunoreactive for Iba1. The integrated intensity of each marker was measured across the master mask (human) or Iba1 mask (mouse) and normalised to the ROI area to give a measure of tissue-wide expression.

#### Single-cell microglial functional marker expression analysis

We next identified all microglia within each ROI and quantify the single-cell expression of Iba1, L-ferritin, HLA-DR, CD68, and CD74, as described in our previous reports [57,58]. Briefly, each object within the previously generated master mask or Iba1 mask was considered a cell, and the average intensity of Iba1, L-ferritin, HLA-DR, CD68, and/or CD74 was measured in each cell. The average intensity of each marker per cell is equivalent to the protein concentration per cell.

Based on the average intensity, we categorised each cell as either high- or low-expressing for each marker of interest (MOI). Distinct low- and high-expressing microglial populations were only identified for Iba1 (Fig. 3M). To determine low/high thresholds for the remaining four functional markers, all microglia from all cases were first plotted on an *XY* scatter plot for the average intensity of Iba1 and each other functional marker, as previously described [57,58] (Fig. S4). The distribution of cells on these *XY* scatter plots was used to determine thresholds for the remaining functional markers.

#### T-SNE plot visualisation and cluster analysis

The t-SNE plots were generated in R using the Barnes-Hut t-SNE implementation package ‘Rtsne v.0.15’ [59,60]. To ensure equal sampling from each region and case, we randomly subsampled 31,200 cells from each case/region subgroup (control motor cortex, ALS motor cortex, control hippocampus, and ALS hippocampus) for a total of 156,000 cells. Single-cell average intensities of L-ferritin, HLA-DR, CD68, CD74, and Iba1 were log transformed for equal weighting and selected for t-SNE dimensional reduction. Parameters were set using Kobak and Berens (2019), Belkina et al. (2019), and Cao et al. (2019) as guidelines [61–63]. Briefly, perplexity values between 30 and 250 were utilized with 250 found to be more visually appealing. A learning rate of 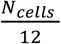 was set where 12 corresponds to the early exaggeration (EE) factor. The EE factor was applied for the first 250 iterations of the maximum 1000 conducted.

Clusters were generated on the 2D t-SNE embedding using a KNN (k nearest neighbour) and Louvain clustering approach in R with the ‘FNN v.1.1.3’ and ‘igraph v.1.2.11’ packages [64,65]. The k nearest neighbour value was set as 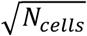 and the NN distances generated were used in the Louvain clustering algorithm to obtain the 27 total cluster assignments. Using these cluster assignments, we calculated the contribution of control versus ALS microglia to each cluster, the percentage of microglia from a given case type in each cluster (done separately for the motor cortex and hippocampus), and the percentage of MOI^high^ microglia in each cluster.

### Statistical analyses

For human analyses, microglial densities, microglial area load, pathology loads, MOI tissue-wide integrated intensities, mean single-cell MOI average intensities, and MOI^high^ percentages were compared between case groups with multiple Mann-Whitney tests and multiple comparisons were controlled for using a False Discovery Rate of 0.01, as determined by the two-stage step-up method of Benjamini, Krieger, and Yekutieli. Statistical significance was set at p ≤ 0.05 with significance of differences between case groups shown as ****p ≤ 0.0001, *** p ≤ 0.001, **p ≤ 0.01, *p ≤ 0.05. Load of pTDP-43 and GFAP, the tissue-wide MOI integrated intensities, the percentage of L-ferritin^high^, HLA-DR^high^, CD68^high^, CD74^high^, and Iba1^high^ microglia, age, and post-mortem delay were sequentially correlated with one another using Spearman correlations. Statistical significance was set at p ≤ 0.05 and all r values greater than 0.7 were considered very strong correlations.

For mouse analyses, pTDP-43 load, microglial cell density, and the percentages of CD68^high^ and L-ferritin^high^ microglia were compared between groups at each time point with a 2-way ANOVA and multiple comparisons were corrected for using Bonferroni’s multiple comparisons test. Statistical significance was set at p ≤ 0.05 with the significance of differences between groups shown as *p ≤ 0.05, ** p ≤ 0.01, ***p ≤ 0.001, ****p≤ 0.0001.

## Results

### Microglia express functional markers in control and ALS post-mortem human motor cortex and hippocampus

To investigate microglial changes in the post-mortem human ALS brain, we developed a 2-round multiplex immunohistochemistry panel to label microglia, ALS pathology, and anatomy in the motor cortex (Fig. 1A-H) and hippocampus (Fig. 1I-P). Round 1 was comprised of microglial markers; L-ferritin, HLA-DR, CD68, CD74, and Iba1 (Fig. 1A-B, E-F, I-J, and M-N). We chose these microglial markers as they have been the most used in previous human ALS immunohistochemistry studies and report on microglial functions (Table S1). Round 2 was comprised of ALS pathology and anatomical markers; pTDP-43, neuronal marker NeuN, astrocyte marker GFAP, and blood vessel marker lectin (Fig. 1C-D, G-H, K-L, and O-P). These rounds were optimised to minimise bleed-through between adjacent channels or antibody cross-reactivities, either within the same round or between rounds after stripping (Fig. S1).

**Figure 1:**
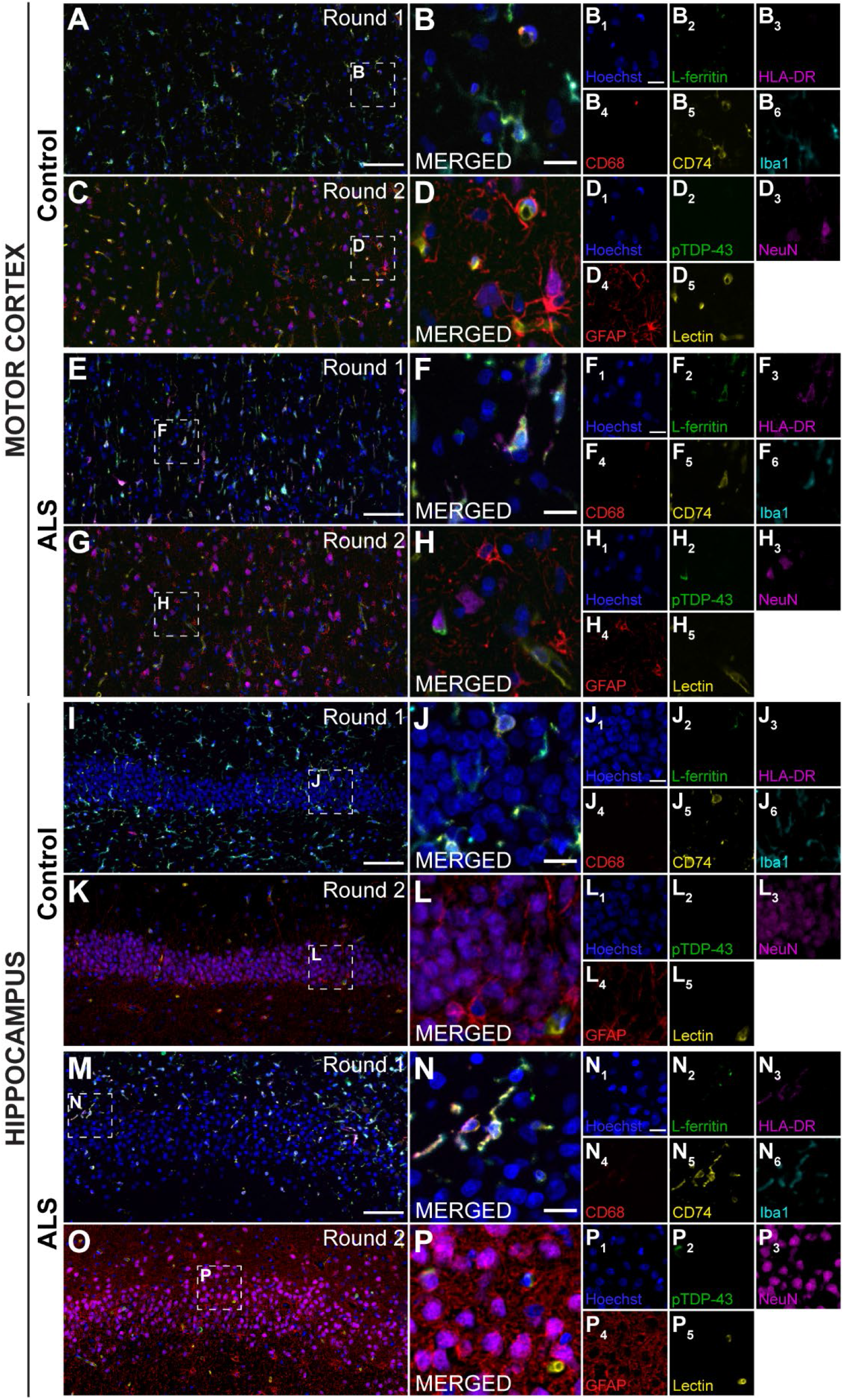
Microglia express functional markers in the normal and ALS post-mortem human motor cortex and hippocampus. Two rounds of immunohistochemical labelling were used to visualise microglial, anatomical, and pathological markers in the motor cortex (**A-H**) and hippocampus (**I-P**) from human ALS (stage 4 shown) and neurologically normal control cases. Round 1 was comprised of microglial markers, including L-ferritin (green), HLA-DR (magenta), CD68 (red), CD74 (yellow), and Iba1 (cyan), with a Hoechst nuclear counterstain (blue) (**A-B, E-F, I-J, M-N**). Round 2 was comprised of pathological and anatomical markers, including pTDP-43 (green), NeuN (magenta), GFAP (red), and lectin (yellow) with a Hoechst nuclear counterstain (blue) (**C-D, G-H, K-L, O-P**). Images from rounds 1 and 2 were aligned using the Hoechst nuclear counterstain. Scale bars (**A, C, E, I, K, M, O**) = 100 μm and scale bars (**B, D, F, H, J, L, N, P**) = 20 μm.

In the first round of labelling, immunoreactivity for all microglial markers was observed in both the ALS and control motor cortex and hippocampus. Of these previously known myeloid-specific markers, HLA-DR, CD68, and CD74 were exclusively expressed by Iba1-positive microglia. In contrast, while L-ferritin has also been shown to be specifically expressed by microglia [30], L-ferritin expression was observed in both Iba1-expressing microglia and GFAP-expressing astrocytes in the motor cortex and hippocampus of ALS and control cases (Fig. S4). As such, L-ferritin immunoreactivity was not used to identify microglia in the quantitative analyses, but microglial L-ferritin immunoreactivity was measured in microglia identified using the other four microglial-specific markers. Of the second round of labelling, NeuN, GFAP, and lectin were observed in both the ALS and control motor cortex and hippocampus. In contrast, pTDP-43 immunoreactivity in the motor cortex and hippocampus distinguished control and ALS cases as previously described [50]; in control cases, no pTDP-43 immunoreactivity was observed in the motor cortex or hippocampus (Fig. 1D_2_ and L_2_), in stage 1-3 ALS, pTDP-43 immunoreactivity was observed in the motor cortex but not the hippocampus, and in stage 4 ALS, pTDP-43 immunoreactivity was observed in both the motor cortex and hippocampus (Fig. 1H_2_ and P_2_).

### Microglial cell density, pTDP-43 load, and GFAP load are increased in human ALS

We next investigated whether there are microglial density changes in ALS and whether microglial density spatially correlates with pTDP-43 aggregate load or astrogliosis (Fig. 2). To do this, we quantified microglial density (Fig. 2A-B), pTDP-43 integrated intensity (Fig. 2C-D), and GFAP area coverage (Fig. 2E-F) in the motor cortex and hippocampus of control and ALS cases.

**Figure 2:**
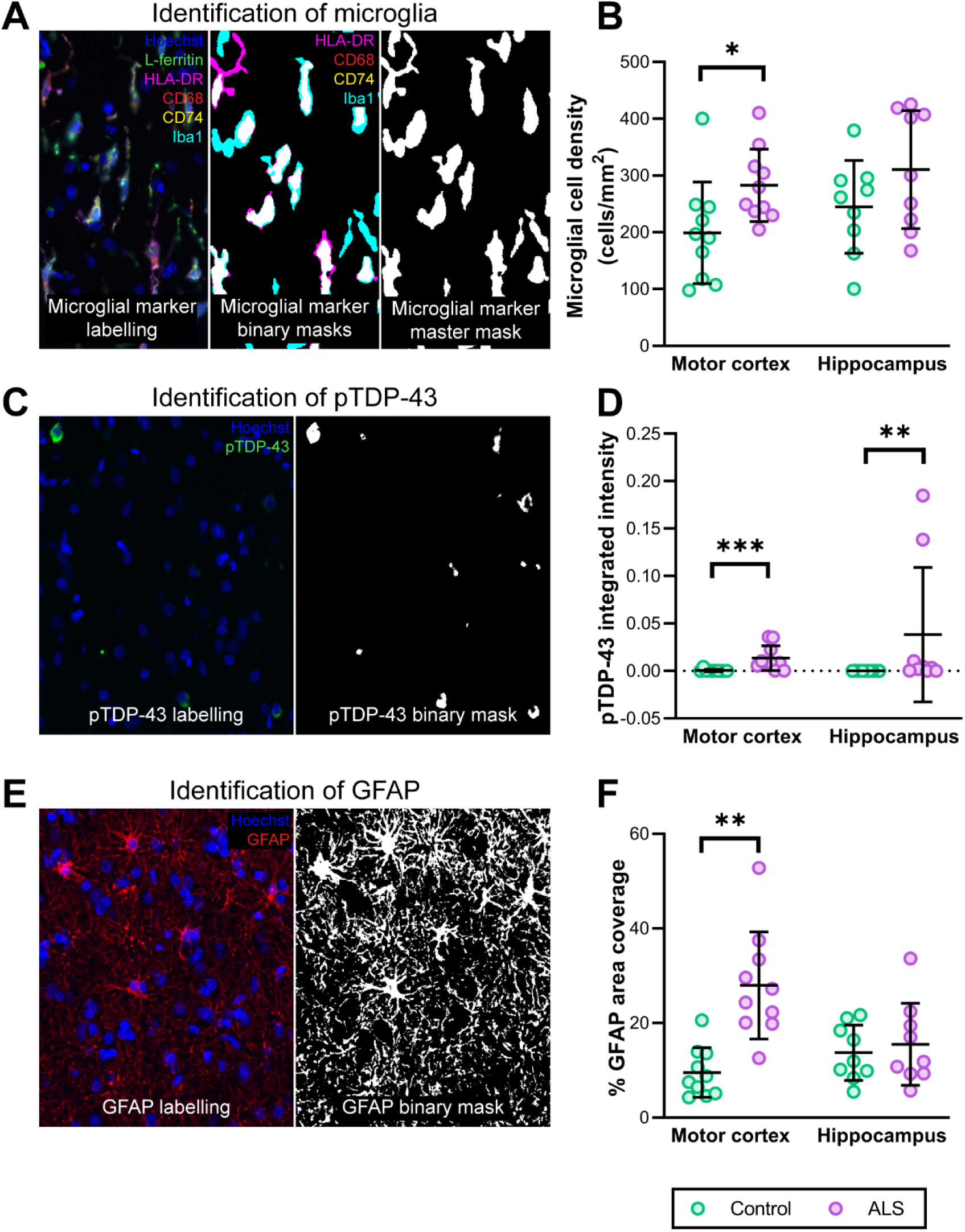
Microglial cell density, pTDP-43, and GFAP load are increased in ALS. Microglial cell density was quantified from round 1 immunolabelling (**A** and **B**) and pTDP-43 load and astrogliosis was quantified from round 2 (**C-F**). Total microglia were identified by creating separate microglial marker binary masks from HLA-DR, CD68, CD74, and Iba1, which were then combined to create a microglial marker master mask (**A**). Each object within this mask was considered a single microglial cell. The microglial cell density in the motor cortex and hippocampus was quantified and compared between control and ALS cases (**B**). Aggregates of pTDP-43 were detected, and binary masks of immunoreactivity were generated (**C**). The integrated intensity of pTDP-43 aggregates identified was quantified and normalised to total tissue are to generate a load measure. The pTDP-43 load was compared between control and ALS motor cortex and hippocampus (**D**). The area of GFAP immunoreactivity identified was quantified and normalised to total tissue are to generate a measure of astrogliosis (**E** and **F**). Data presented as mean ± SD; control n = 10 and ALS n = 9-10. Microglial densities and pathology loads were compared between case groups with multiple Mann-Whitney tests and multiple comparisons were controlled for using a False Discovery Rate of 0.01, as determined by the two-stage step-up method of Benjamini, Krieger, and Yekutieli. Significance of differences between case groups: ****p ≤ 0.0001, **p ≤ 0.01, *p ≤ 0.05.

To quantify microglial density, all microglia were identified by generating single binary masks from HLA-DR, CD68, CD74, or Iba1 immunoreactivity, and these binary masks were combined to generate a master mask of all microglia (Fig. 2A). The total microglial density (where each object within this master mask was considered a single cell) was increased in the ALS motor cortex relative to controls (Fig. 2B, p = 0.0147). In contrast, microglial cell density was unchanged in the ALS hippocampus when compared with controls (Fig. 2B, p = 0.222).

To quantify pTDP-43 aggregate load, we measured the integrated intensity of pTDP-43 immunoreactivity (Fig. 2C). In accordance with presence of pTDP-43 aggregates in the motor cortex being discriminatory between ALS and control cases, the pTDP-43 integrated intensity across the six layers of the motor cortex was significantly increased in ALS relative to control cases (Fig. 2D, p = 0.000487). Furthermore, pTDP-43 integrated intensity was also significantly increased in ALS hippocampus relative to controls (Fig. 2D, p = 0.00276). It is important to note that this significant increase in the hippocampus was driven by stage 4 ALS cases, specifically cases MN15 and MN30 (Table 1).

To measure astrogliosis, we quantified the tissue area occupied by GFAP immunoreactivity (Fig. 2E). We observed a significant increase in GFAP area coverage in the ALS motor cortex relative to control cases (Fig. 2F, p = 0.000206). In contrast, GFAP area coverage was not different between the ALS hippocampus relative to control cases (Fig. 2F, p = 0.796).

Therefore, significant increases in pTDP-43 load were observed in both the human ALS motor cortex and hippocampus. In contrast, increases in microglial cell density and astrogliosis were confined to the ALS motor cortex.

### Microglia increase expression of CD68 and Iba1 in the human ALS motor cortex, but not hippocampus

The significant increase in microglial density in ALS demonstrates microglial changes are occurring in disease. However, an increase in microglial cell number is only one aspect of the microglial response to damage and disease [66]. Therefore, we hypothesised that microglia may also up-regulate specific functional markers in response to ALS neurodegeneration. To investigate this, we quantified the levels of L-ferritin (Fig. 3A-C), HLA-DR (Fig. 3D-F), CD68 (Fig. 3G-I), CD74 (Fig. 3J-L), and Iba1 (Fig. 3M-O) in the ALS and control motor cortex and hippocampus. To quantify the levels of each functional marker, we measured tissue-wide and single-cell immunolabelling intensities using novel automated image analysis pipelines (Fig. S2).

**Figure 3:**
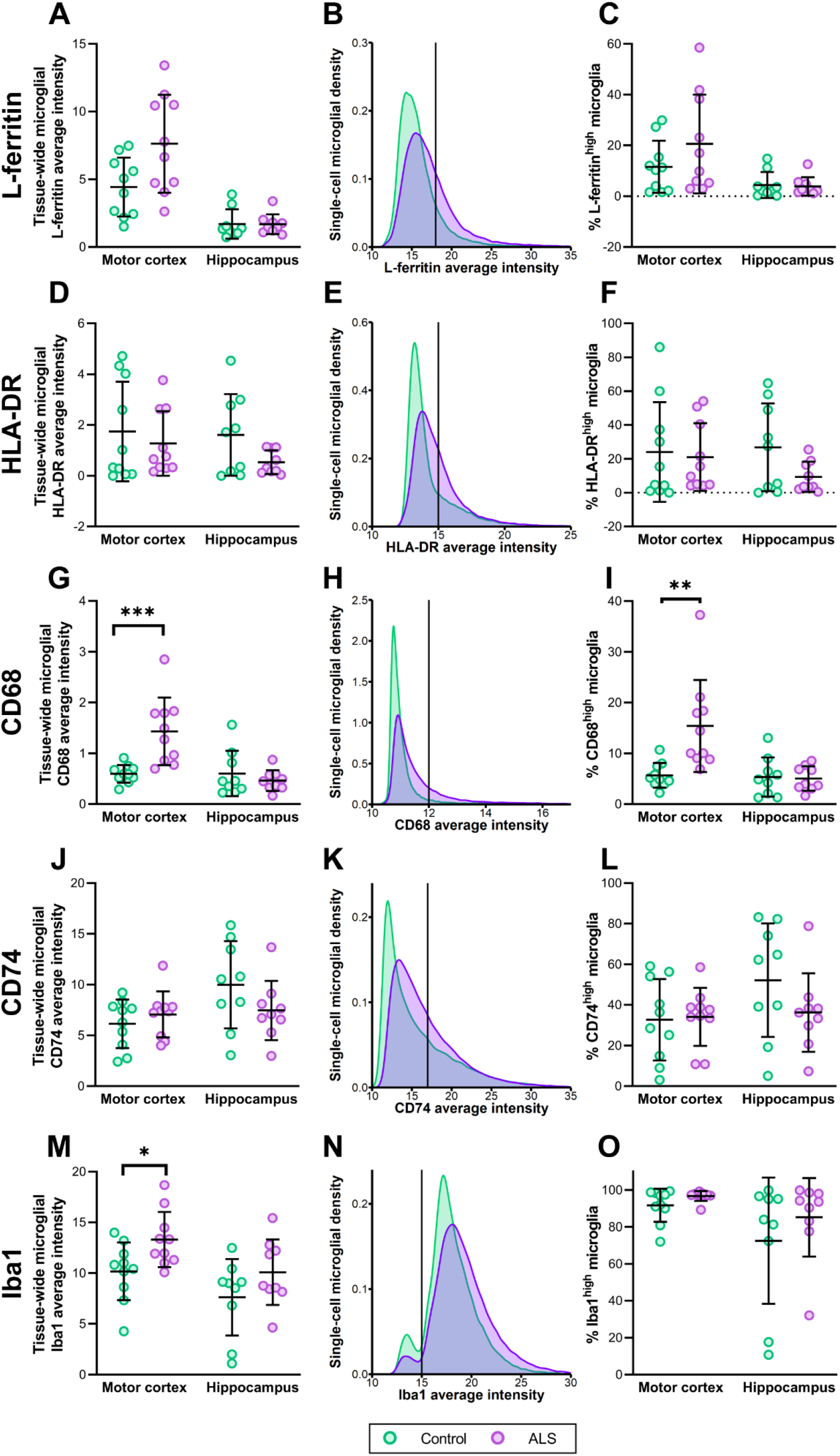
Microglia increase expression of CD68 and Iba1 in the ALS motor cortex. Microglia were identified by combined labelling of all microglial markers from round 1 and the tissue-wide and single cell average intensities of each of these markers was measured and used to quantify changes in functional marker expression in ALS; L-ferritin (**A-C**), HLA-DR (**D-F**), CD68 (**G-I**), CD74 (**J-L**), and Iba1 (**M-O**). Immunolabelling intensity of each functional marker was measured within the microglial master mask and normalised to the area of the microglial master mask. The tissue-wide average intensities were compared between control and ALS cases in the motor cortex and hippocampus for each functional marker (**A, D, G, J, and M**). Data presented as mean ± SD; control n = 10 and ALS n = 9-10. To designate cells as either high- or low-expressing for each functional marker of interest (MOI^high^ or MOI^low^), total microglia from all control and all ALS cases were pooled and the distribution curve for each marker was generated (**B, E, H, K, and N**). The threshold for each marker was determined and each cell was designated as either MOI^high^ or MOI^low^ for each functional marker. The percentage of MOI^high^ cells was compared between control and ALS cases in the motor cortex and hippocampus for each functional marker (C, F, I, L, and O). Data presented as mean ± SD; control n = 10 and ALS n = 9-10. MOI average intensities and MOI^high^ percentages were compared between case groups with multiple Mann-Whitney tests and multiple comparisons were controlled for using a False Discovery Rate of 0.01, as determined by the two-stage step-up method of Benjamini, Krieger, and Yekutieli. Significance of differences between case groups: *p ≤ 0.05, ** p ≤ 0.01, ***p ≤ 0.001.

We first investigated whether there was a tissue-wide increase in microglial expression of each functional marker (Fig. 3A, D, G, J, and M). We quantified the average intensity of immunofluorescent labelling for each functional marker within the microglial master mask, equivalent to the average expression of each marker across all microglia in each case. The average intensity of each functional marker was compared in the motor cortex and hippocampus between control and ALS cases. We identified significant increases in CD68 and Iba1 in the ALS motor cortex (CD68: p = 0.000206, Fig. 3G; Iba1: p = 0.0185, Fig. 3M). All other markers were unchanged in the ALS motor cortex, while all functional markers were unchanged in the ALS hippocampus (Fig. 3).

We hypothesised that the tissue-wide increases in microglial CD68 and Iba1 expression were due to an increase in the number of high-expressing microglia. Based on the single-cell average intensity of each functional marker and a manually determined threshold, we classified each microglia as either high or low expressing for each marker (Fig. 3B, E, H, K, and N). The percentage of cells expressing the marker of interest highly (MOI^high^) was compared between control and ALS cases for each functional marker (Fig. 3C, F, I, L, and O). In accordance with the significant increase in the tissue-wide expression of CD68 in the ALS motor cortex, we identified a significant increase in the percentage of CD68^high^ cells in the ALS motor cortex when compared with controls (p = 0.000444, Fig. 3I). In contrast, despite the tissue-wide increase in Iba1 levels, the percentage of Iba1^high^ microglia was not significantly different in the ALS motor cortex from controls (p = 0.424, Fig. 3O), likely because nearly all microglia had high levels of Iba1 in the control brain. We did not identify any change in other functional markers in the ALS motor cortex, and no functional markers were increased in the ALS hippocampus (Fig. 3C, F, I, L and O).

Therefore, CD68 expression is robustly increased in ALS and shows both a tissue-wide increase in expression and an increase in CD68^high^ microglia. In contrast, Iba1 expression is significantly increased in the ALS motor cortex due to already high-expressing microglia further increasing their Iba1 expression.

### CD68 expression correlates with pTDP-43 pathology load in the ALS motor cortex

We next determined whether the increase in the percentage of CD68^high^ microglia correlated with the expression of other microglia functional markers or measures of ALS neurodegeneration (Fig. 4). Therefore, measures of pTDP-43 integrated intensity and GFAP area coverage, and the percentages of L-ferritin^high^, HLA-DR^high^, CD68^high^, CD74^high^, and Iba1^high^ microglia were sequentially correlated with one another within the control and ALS motor cortex and hippocampus.

**Figure 4:**
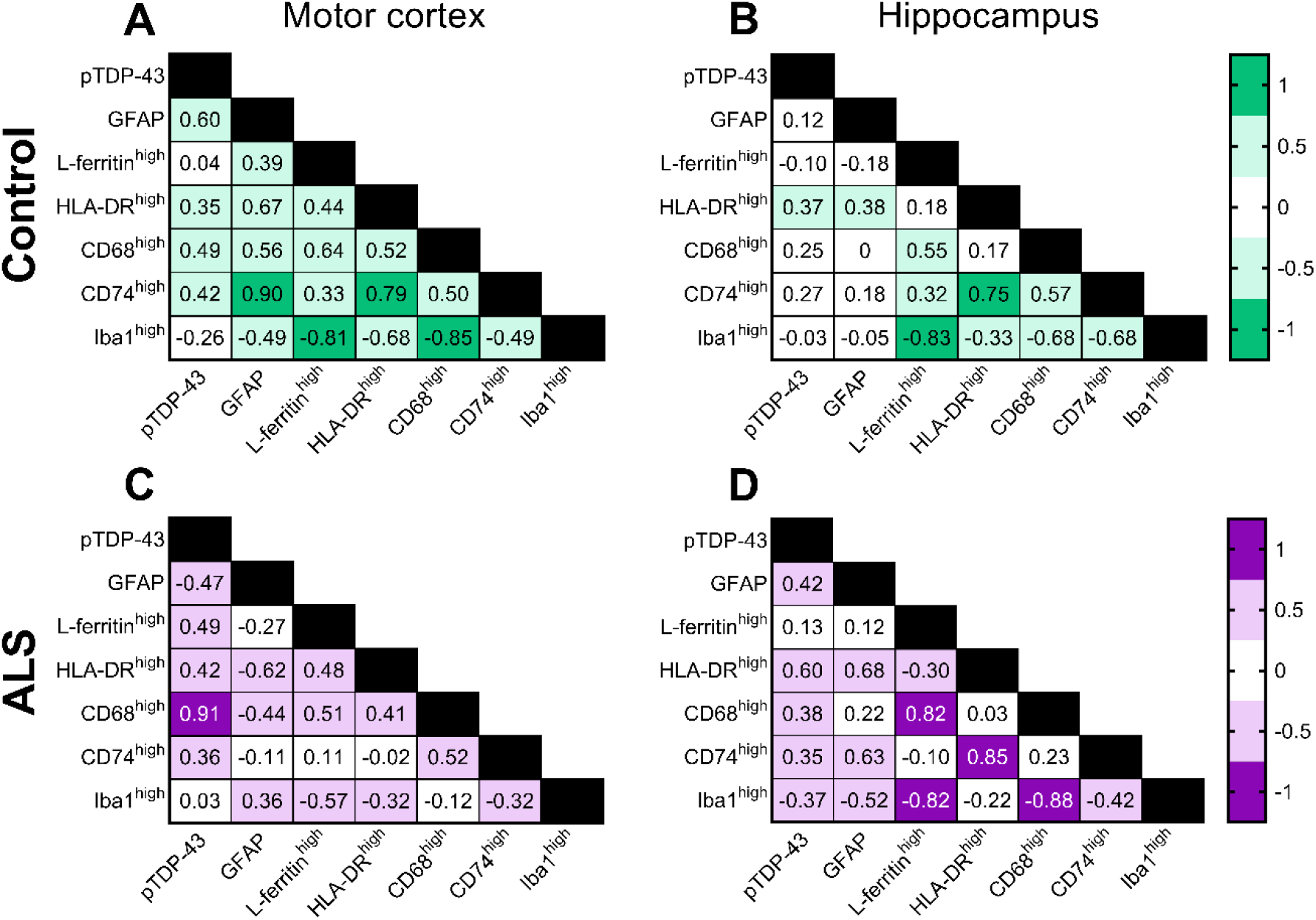
Percentage of CD68^high^ microglia correlates with pTDP-43 load in the ALS motor cortex. Measures of microglial functional changes and neuropathology load were correlated in the control (**A-B**) and ALS (**C-D**) motor cortex and hippocampus. Measures of pTDP-43 integrated intensity and GFAP area coverage, and percentages of L-ferritin^high^, HLA-DR^high^, CD68^high^, CD74^high^, and Iba1^high^ microglia were sequentially correlated with one another using Spearman correlations. The resulting r value from each correlation is presented in the correlation matrix and colour coded relative to strength. All r values greater than 0.7 were statistically significant and considered very strong correlations. Control n = 10 and ALS n = 9-10, where statistical significance was set at p ≤ 0.05.

Co-expression of certain microglial functional markers was expected even in the normal human brain [30], so it was first important to first ascertain the relationships between microglial markers and measures of neurodegeneration in control cases. In the control motor cortex, strong or very strong statistically significant linear correlations were observed between the percentages of CD74^high^ and HLA-DR^high^ microglia (r = 0.794, p = 0.00880), Iba1^high^ and L-ferritin^high^ microglia (r = −0.806, p = 0.00720), and Iba1^high^ and CD68^high^ microglia (r = −0.855, p = 0.00290), as well as the percentage of CD74^high^ microglia and GFAP area coverage (r = 0.903, p = 0.000800), and the percentage of HLA-DR^high^ microglia and GFAP area coverage (r = 0.673, p = 0.0390; Fig. 4A). A similar pattern of correlations was observed in the control hippocampus, with strong or very strong statistically significant correlations identified between the percentages of CD74^high^ and HLA-DR^high^ microglia (r = 0.750, p = 0.0255), Iba1^high^ and L-ferritin^high^ microglia (r = −0.833, p = 0.00830), Iba1^high^ and CD68^high^ microglia (r = −0.683, p = 0.0503), and Iba1^high^ and CD74^high^ microglia (r = −0.683, p = 0.0503), although the latter did not reach statistical significance (Fig. 4B).

We subsequently carried out the same sequence of Spearman correlations between the percentages of MOI^high^ microglia and pathology loads in the ALS motor cortex and hippocampus (Fig. 4C and D). The pattern of correlations in ALS was compared with control cases to determine 1) the emergence of unique correlations between pairs of microglial markers and 2) the loss of baseline normal correlations. The pattern of correlations in the ALS hippocampus mirrored that of the control motor cortex and hippocampus; strong or very strong statistically significant correlations were observed between the percentages of CD68^high^ and L-ferritin^high^ microglia (r = 0.817, p = 0.0108), CD74^high^ and HLA-DR^high^ microglia (r = 0.850, p = 0.00610), Iba1^high^ and L-ferritin^high^ microglia (r = −0.817, p = 0.0108), Iba1^high^ CD68^high^ microglia (r = −0.883, p = 0.00310), and HLA-DR^high^ microglia and GFAP load (r = 0.683, p = 0.0503), although this did not reach statistical significance. In contrast, pattern of correlations in the ALS motor cortex was unique. No statistically significant correlations were identified between the percentages of MOI^high^ microglia. Instead, a very strong statistically significant correlation was identified between the percentage of CD68^high^ microglia and pTDP-43 load (r = 0.906, p = 0.0006). A similar pattern of correlations was observed when using the tissue-wide integrated intensity measures in the place of MOI^high^ percentages (Fig. S5). We did not identify any notable correlations between pathology load or MOI^high^ percentages with age or post-mortem delay in either the ALS or control case groups (Table S2).

### Microglial subpopulations that are increased in ALS motor cortex are defined by high L-ferritin levels

Given microglial heterogeneity in the human brain, we hypothesised that changes in single functional proteins would not reflect the complexity of ALS-associated microglial changes. Our multiplex immunohistochemistry and image analysis pipelines allowed for the measurement of five functional proteins by each microglia, creating a multi-dimensional dataset from post-mortem tissues.

K-Nearest neighbour distancing in conjunction with Louvain clustering on the t-SNE embedding allowed for the identification of 27 clusters within the data. However, none of the 27 microglial clusters identified were spatially distinct within the t-SNE plots (Fig. 5A). We attributed this to the low dimensionality of our dataset; microglia were clustered based on the single-cell abundance of 5 functional proteins expressed in a narrow physiological range. The 27 microglial clusters identified therefore do not represent unique microglial subpopulations in the control and ALS brains. Instead, each cluster represents a subset of microglia with subtle differences in expression of L-ferritin, HLA-DR, CD68, CD74, and/or Iba1. Therefore, in the context of this study, we quantified changes in the abundance of the microglial clusters identified to signify subtle changes in microglial functional protein levels that would have been missed by tissue-wide analyses.

**Figure 5:**
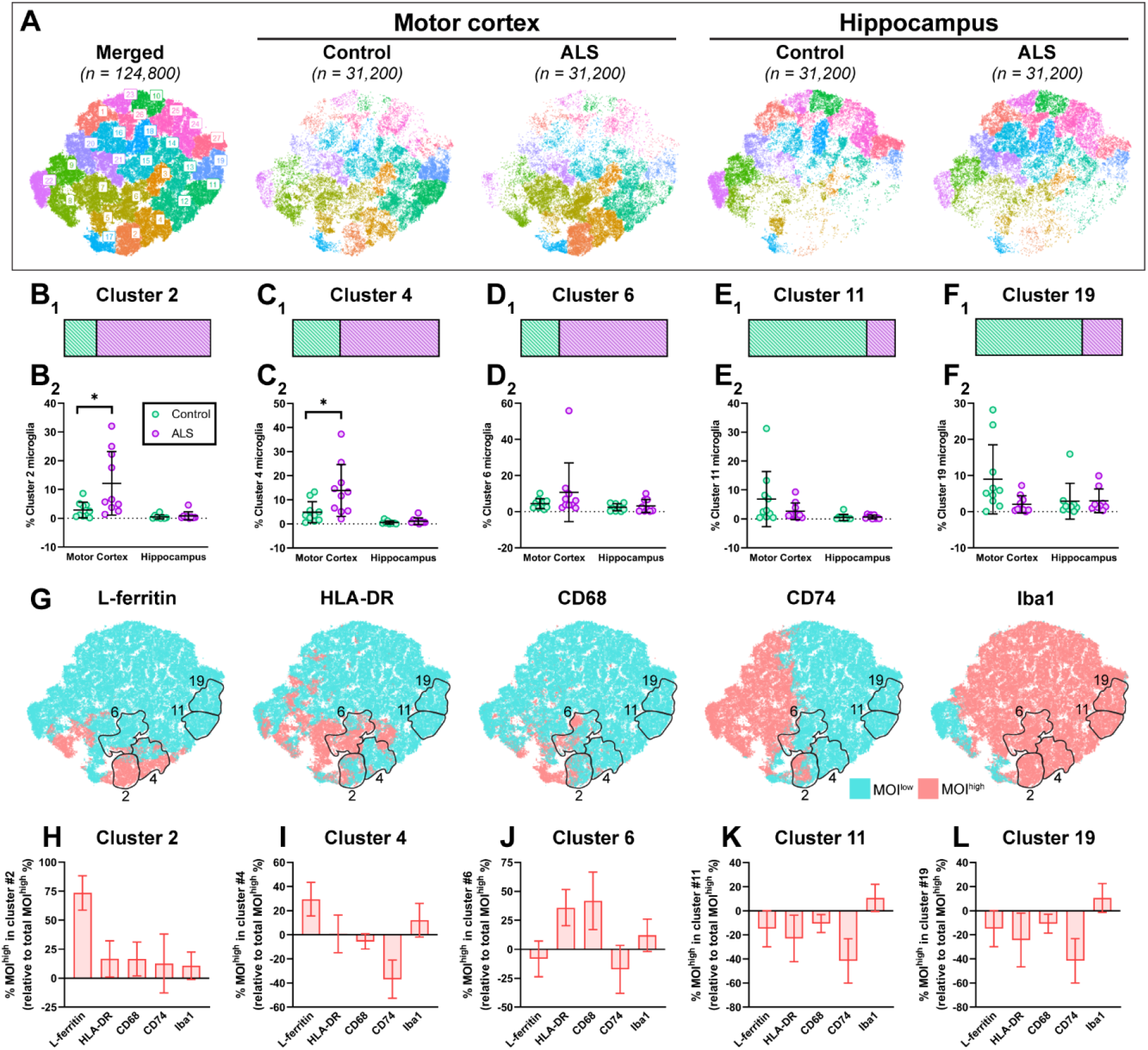
Microglial clusters enriched ALS motor cortex express high levels of L-ferritin. Microglial single-cell average intensities of L-ferritin, HLA-DR, CD68, CD74, and Iba1 were used to identify and characterise microglial subpopulations that were changed in ALS relative to controls. A t-SNE plot was generated using random subset of 124,800 microglia (31,200 each from control motor cortex, ALS motor cortex, control hippocampus, and ALS hippocampus), and 27 clusters were identified using a k nearest neighbour and Louvain clustering approach (**A**). The total contribution of control versus ALS microglia to each cluster was visualised with a percentage bar to determine clusters most different between control and ALS: clusters 2, 4, 6, 11, and 19 (**B_1_-F_1_**). The percentage of total microglia in clusters 2, 4, 6, 11, and 19 were statistically compared between control and ALS motor cortex and hippocampus (**B_2_-F_2_**). Data presented as mean ± SD; control n = 10 and ALS n = 9-10. Cluster percentages were compared between case groups with multiple Mann-Whitney tests and multiple comparisons were controlled for using a False Discovery Rate of 0.01, as determined by the two-stage step-up method of Benjamini, Krieger, and Yekutieli. Significance of differences between case groups: *p ≤ 0.05. Microglial single-cell average intensities of L-ferritin, HLA-DR, CD68, CD74, and Iba1 were mapped on the merged t-SNE plot (all microglia from ALS and control motor cortex and hippocampus) using a binary high-low approach to visualise MOI^low^ and MOI^high^ microglia (G). To phenotype clusters based on MOI^high^ percentage, for each functional marker the percentage of MOI^high^ microglia in each cluster for a given case to the percentage of MOI^high^ microglia in that case overallThe percentage MOI^high^ for each functional marker was normalised to the total percentage of MOI^high^ in each case for clusters 2, 4, 6, 11, and 19 in all cases (H-L). Data presented as mean ± SD; n = 20.

None of the 27 microglial clusters identified were unique to either the control or ALS brain. However, within both case types, we noted anatomical regional patterning; 18 clusters were differentially abundant in the motor cortex versus the hippocampus (Fig. 5A, Fig. S6). Therefore, the pattern of coexpression of the 5 functional markers is inherently different between motor cortex and hippocampal microglia irrespective of disease state.

Five clusters (2, 4, 6, 11, 19) were disproportionately contributed to by control or ALS microglia (Fig. 5B_1_-F_1_). For these clusters we then quantified the percentage of all microglia for each case that fell into each cluster and compared these percentages between control and ALS motor cortex and hippocampus (Fig. 5B_2_-F_2_). Microglia that fell into clusters 2 and 4 were significantly enriched in the ALS motor cortex compared with controls (p = 0.0115, Fig. 5B_2_ and C_2_). In contrast, microglia that fell into clusters 6, 11, and 19 were not consistently enriched in either ALS or control cases (Fig 5D_2_-F_2_). Instead, the disproportionate contribution of ALS microglia to cluster 6 (Fig. 5D_1_) and control microglia to clusters 11 and 19 (Fig. 5E_1_-F_1_) were driven by one or two cases, suggesting the distribution of microglia across the t-SNE generated is different in each case. Percentage data for all clusters is presented in Figure S8.

With microglial clusters 2, 4, 6, 11, and 19 being enriched with control or ALS microglia, and ALS cases being enriched with microglia from clusters 2 and 4, albeit with some case variability, we sought to characterise the 5-functional-marker phenotype of each cluster. To visually phenotype clusters 2, 4, 6, 11, and 19, we mapped the single-cell average intensities of L-ferritin, HLA-DR, CD68, CD74, and Iba1 on merged t-SNEs (all microglia from ALS and control motor cortex and hippocampus) as a binary high-low approach to visualise MOI^low^ and MOI^high^ microglia (Fig. 5G).

We then quantified the functional marker phenotype of each microglial cluster (Fig. 5H-L); we normalised the percentage of MOI^high^ microglia that fell within the cluster, for a given case, to the percentage of MOI^high^ microglia in that case overall; a relative MOI^high^ percentage greater than zero therefore denoted a greater percentage of MOI^high^ microglia in the cluster relative to the case overall. As such, we were able to use the relative percentage of MOI^high^ microglia in a cluster to identify the functional markers driving microglial clustering, and to better understand the functional phenotype of microglial clusters; in particular, clusters 2 and 4 that were enriched in the ALS motor cortex. Cluster 2 was classified by a high relative percentage of L-ferritin^high^ CD68^high^ microglia, cluster 4 by a high relative percentage of L-ferritin^high^ CD74^low^ microglia, cluster 6 by a high relative percentage of HLA-DR^high^ CD68^high^ microglia, and clusters 11 and 19 by high relative percentages of HLA-DR^low^ CD68^low^ CD74^low^ microglia.

With the significant increase in the percentage of CD68^high^ microglia in the ALS motor cortex, we expected an enrichment of microglial clusters defined by high CD68 expression in ALS. However, microglial clusters characterised by high CD68 expression included clusters 1, 2, 5, 6, 8, and 17, of which only cluster 2 was significantly enriched in the ALS motor cortex (Fig. S7 and S8). Therefore, the increase in CD68 expression can be accompanied by either up- or down-regulation of other microglial functional markers.

In summary, CD68^high^ microglia were distributed across many clusters and were particularly abundant in cluster 2 which, along with cluster 4, was enriched in the ALS motor cortex and characterised by a high level of L-ferritin expression.

### pTDP-43-dependent increase in CD68 and L-ferritin in a TDP-43-driven mouse model of ALS

End-stage disease post-mortem human tissue cannot provide temporal information on changes throughout disease. While we have identified changes in microglial CD68 and L-ferritin expression in the human ALS brain, it is unclear <u>when</u> these changes occur relative to pTDP-43 aggregation and neuronal degeneration. The strong correlation between the percentage of CD68^high^ microglia and pTDP-43 load in the ALS motor cortex suggests a relationship between pTDP-43 aggregation and a change in microglial function. To understand this relationship temporally, we utilised brain tissue from bigenic *NEFH*-tTA/*tetO*-hTDP-43ΔNLS (rNLS) and single transgenic *tetO*-hTDP-43ΔNLS (control) mice at 2, 4, and 6 weeks off doxycycline (WOD), equivalent to disease onset, early disease, and late disease stages, respectively [51].

We immunolabelled the rNLS and control mouse brain sections with microglial markers CD68, L-ferritin, and Iba1, pTDP-43, and neuronal marker NeuN (Fig. 6A-J). All microglial markers were observed in both the rNLS and control motor cortex and hippocampus at 2, 4, and 6 WOD. As in the human brain tissue, CD68 was exclusively expressed by Iba1-positive microglia (Fig. 6A-J). L-ferritin expression was observed by Iba1-positive microglia, however, as in the human tissue, L-ferritin expression was not specific to Iba1-positive microglia; L-ferritin immunoreactivity was also observed in other cell types not identified by this panel. Immunoreactivity for pTDP-43 was exclusively identified in rNLS mice, both in the motor cortex and hippocampus (Fig. 6B_5_, D_5_, G, and J).

**Figure 6:**
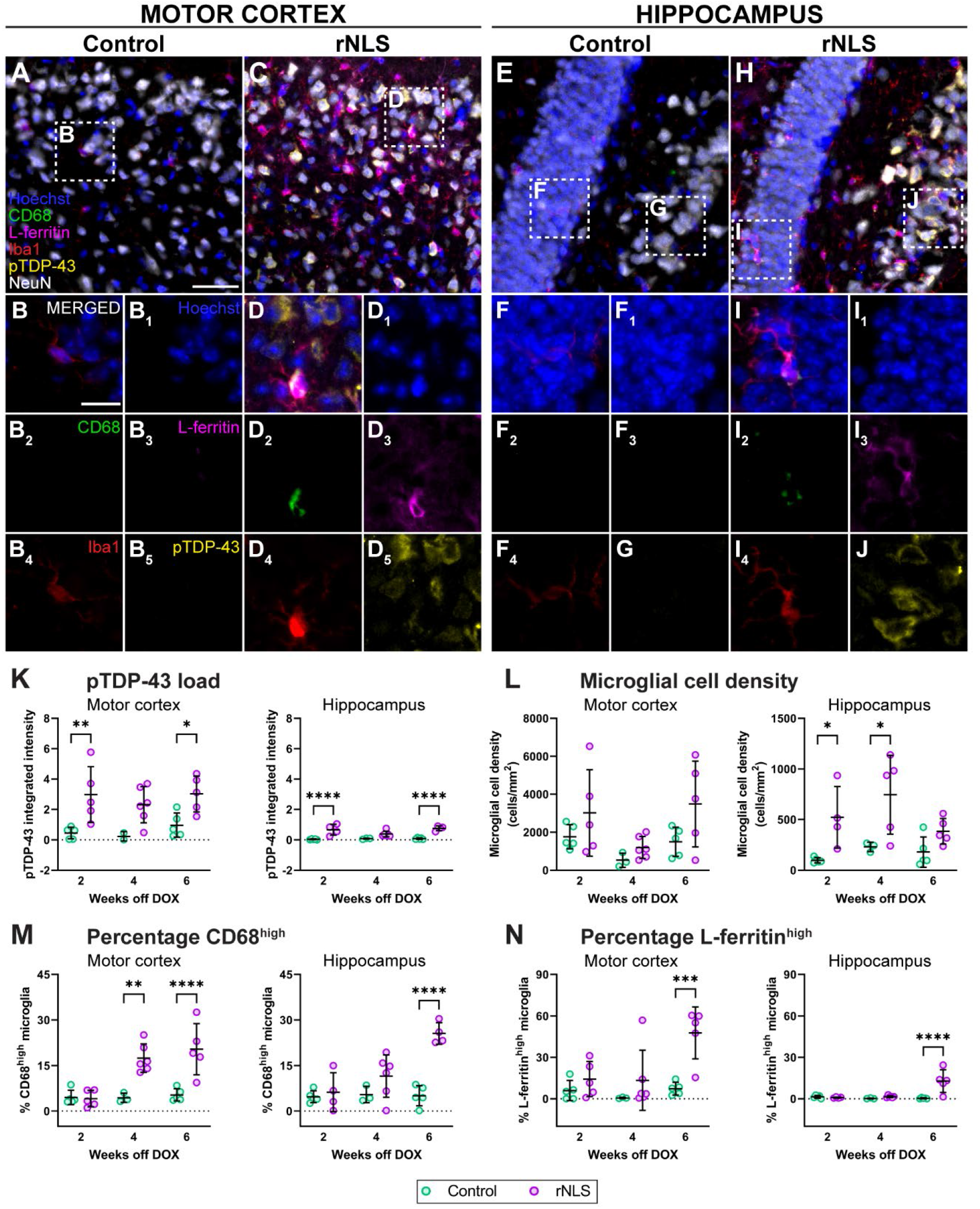
Microglial CD68 and L-ferritin expression follow pTDP-43 aggregation in motor cortex and hippocampus of rNLS mice. Immunohistochemical labelling was used to visualise microglial, anatomical, and pathological markers in the of bigenic *NFFH*-tTA/*teyO*-hTDP-43ΔNLS (rNLS) and single transgenic *tetO*-hTDP-43ΔNLS (control) mice at 2, 4, and 6 weeks off DOX. Example images of the motor cortex (**A-D**) and hippocampus (**E-J**) from mice at 6 weeks of DOX are shown; microglia were labelled with CD68 (green), L-ferritin (magenta), and Iba1 (red), pathology with pTDP-43 (yellow), and neurons with NeuN (white), with a Hoechst nuclear counterstain (blue); scale bar (**A**) = 100 μm and scale bar (**B**) = 20 μm. Microglial cell density (**K**), pTDP-43 load (**L**), and the percentage of CD68^high^ (**M**) and L-ferritin^high^ (**N**) microglia were quantified in the hippocampus and motor cortex of control and rNLS mice 2, 4, and 6 weeks off DOX (K). All data presented as mean ± SD; control n = 4-6 and rNLS n = 2-5. Measurements were compared between case groups at each time point with a 2-way ANOVA and multiple comparisons were corrected for using Bonferroni’s multiple comparisons test. Significance of differences between case groups: *p ≤ 0.05, ** p ≤ 0.01, ***p ≤ 0.001, ****p≤ 0.0001.

Having verified functional marker expression, we next quantified the pTDP-43 pathology load in the rNLS mice relative to controls at each time point (Fig. 6K). As with the human tissue, to measure pTDP-43 aggregate load, we measured the integrated intensity of pTDP-43 immunoreactivity. At 2 WOD, the pTDP-43 load was significantly higher in rNLS mice relative to controls in both the motor cortex and the hippocampus (motor cortex p = 0.0093, hippocampus p = 0.0001; Fig. 6K), with the pTDP-43 load higher in the motor cortex than hippocampus. This increased pTDP-43 load was maintained in rNLS mice through to 6 WOD in both regions (motor cortex p = 0.0399, hippocampus p < 0.0001; Fig. 6K), although did not reach statistical significance at 4 WOD (motor cortex p = 0.349, hippocampus p = 0.199; Fig. 6K). Therefore, in line with the previous papers describing the rNLS mouse model of ALS, we observed pTDP-43 pathology in the motor cortex and hippocampus from 2 WOD.

We subsequently characterised microglial changes occurring in the rNLS mouse model relative to pTDP-43 aggregation. First, we quantified microglial cell density as a broad measure of microgliosis. Microglial density remained unchanged in the rNLS motor cortex at all time points but was significantly increased in the rNLS hippocampus at 2 and 4 WOD, but not 6 WOD (hippocampus; 2 WOD p = 0.0292, 4 WOD p = 0.0139, 6 WOD p = 0.407; Fig. 6L). However, it is important to note a high level of variability in the data, particularly in the rNLS groups at all disease timepoints. Therefore, while there were some indications of microgliosis in the rNLS mice, a high level of variability meant we were unable to identify a pattern of changing microglial density with pTDP-43 aggregation.

To further phenotype microglial changes, we quantified the percentage of CD68^high^ and L-ferritin^high^ microglia in the motor cortex and hippocampus of the rNLS and control mice using equivalent methodology to our human tissue analysis (Fig. 6M and N). The percentage of CD68^high^ microglia was unchanged at 2 WOD but significantly increased at 4 and 6 WOD in the rNLS motor cortex and hippocampus relative to controls (motor cortex: 2 WOD p = 0.999, 4 WOD p = 0.0160, 6 WOD p = 0.0003; hippocampus: 2 WOD p > 0.999, 4 WOD p = 0.00550, 6 WOD p < 0.0001; Fig. 6M). Further to this, the percentage of L-ferritin^high^ microglia was unchanged at 2 and 4 WOD but significantly increased at 6 WOD in the rNLS motor cortex and hippocampus relative to controls (motor cortex: 2 WOD p = 0.7276, 4 WOD p = 0.406, 6 WOD p = 0.0005; hippocampus: 2 WOD p = 0.991, 4 WOD p = 0.935, 6 WOD p < 0.0001; Fig. 6N). Therefore, while pTDP-43 load was significantly increased at from 2 WOD, the percentage of CD68^high^ and L-ferritin^high^ microglia were increased later, from 4 and 6 WOD, respectively.

## Discussion

In this study, we utilised multiplexed immunohistochemistry and single cell analysis pipelines to identify how microglia change spatiotemporally relative to TDP-43 pathology ALS. In the human ALS motor cortex, we identified increased microglial expression of both Iba1 and CD68, the latter of which strongly correlated with increased pTDP-43 load. Further to this, we identified microglial phenotype clusters enriched in the ALS motor cortex that were defined by high L-ferritin expression. In the rNLS mouse-driven model of ALS, higher microglial CD68 and L-ferritin expression occurred at early and late disease stages, respectively, <u>after</u> pTDP-43 aggregation. The results of this study are summarised in Figure 7. Overall, this work provides a detailed examination of microglial changes in the human ALS brain and their association with pTDP-43 pathology.

**Figure 7:**
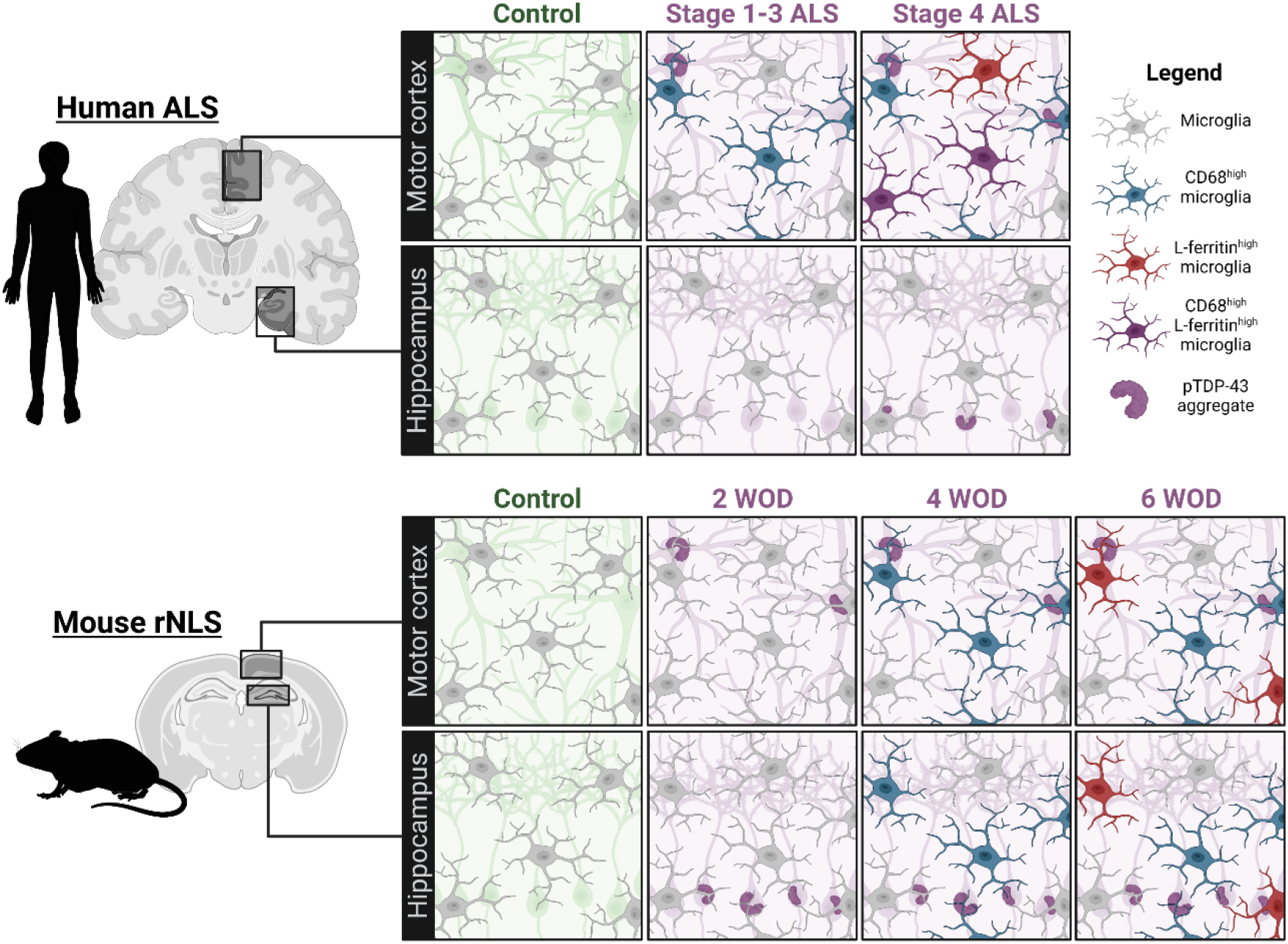
Summary of microglial changes identified in the human ALS and mouse rNLS motor cortex and hippocampus. The human ALS motor cortex showed increased microglial expression of both Iba1 and CD68, the latter of which strongly correlated with increased pTDP-43 load. Microglial phenotype clusters that were enriched in the ALS motor cortex were defined by high L-ferritin expression. In the rNLS mouse-driven model of ALS, higher microglial CD68 and L-ferritin expression occurred at early and late disease stages, respectively, after pTDP-43 aggregation.

We first investigated the levels of specific markers whose expression reflects different microglial function; L-ferritin, HLA-DR, CD68, CD74, and Iba1 (Table S1). The most notable change identified was an increase in CD68, evidenced by increased tissue-wide CD68 average intensity and percentage of CD68^high^ microglia in the human ALS motor cortex, and increased percentage of CD68^high^ microglia in the rNLS mouse motor cortex and hippocampus at early and late disease stages. CD68 is a lysosomal protein classically considered a marker of ‘activated’ phagocytic microglia [36,37]. The robust increase in CD68 expression in both the human ALS motor cortex and rNLS brain suggests that at least a subset of microglia have increased phagocytic activity as part of their reaction to disease pathology. Indeed, microglia have the capacity to phagocytose both pTDP-43 and degenerating neurons, likely an as attempt to rescue the brain from further degeneration [18,67,68]. Therefore, the increase in microglial CD68 expression could suggest a neuroprotective function in ALS.

The main strength of this study was the use of multiplexed immunohistochemistry [56] to investigate the expression of five microglial functional markers alongside anatomy and pathology in the human ALS brain. Using these labelling methods with single-cell image analysis pipelines [57], we identified two microglial clusters more abundant in the human ALS motor cortex, both of which were characterised by high L-ferritin. This was supported by the finding of an increased percentage of L-ferritin^high^ microglia in the late-symptomatic disease rNLS motor cortex and hippocampus. L-ferritin is an iron storage protein, up-regulated by microglia to maintain brain iron homeostasis [29,69]. However, high L-ferritin expression is identified in dystrophic microglia in the normal and diseased human brain, suggesting that excess L-ferritin expression is associated with microglial dysfunction [29,30,57,69,70]. L-ferritin expression by CD68-expressing microglia has previously been reported in the deep layers of the ALS motor cortex [24], and the emergence here of a CD68^high^ L-ferritin^high^ microglial cluster in end-stage human ALS brain suggests that a pool of CD68^high^ phagocytic microglia shift in late disease to an L-ferritin^high^ dysfunctional state. This shift to a dysfunctional state could implicate a loss of microglial neuroprotective functions in secondary neuronal degeneration in ALS.

We subsequently investigated relationships between microglial changes and pTDP-43 aggregation. Microglial CD68 levels significantly correlated with pTDP-43 load in the human ALS motor cortex. This data corroborates a previous report of a correlation between CD68 expression and pTDP-43 pathology in the ALS spinal cord [25]. Nolan and colleagues also demonstrated occasional pTDP-43 aggregates within the processes of CD68-positive microglia, proposing that microglial phagocytosis of pTDP-43 aggregates drives CD68 expression [25]. It is also possible that the pTDP-43 within microglial processes is of microglial rather than neuronal origin, however the expression of phagocytosis-related lysosomal marker CD68 suggests that they are acquired by phagocytosis. In support of this, CD68^high^ microglia are also observed in the rNLS mouse model of ALS, in which human TDP-43 lacking a nuclear localisation signal is expressed only within neurons and a small proportion of astrocytes but not in microglia or oligodendrocytes [51]. Furthermore, microglia are capable of phagocytosing TDP-43 of neuronal origin [18,67,68]. Indeed, when microglia lose their capacity to phagocytose TDP-43 through TREM2 depletion, microglia are locked into a homeostatic state [67]. Therefore, while the mechanism driving microglial CD68 expression requires further study, we have provided further evidence of a relationship between microglial phagocytic function and the TDP-43 pathology observed in the human motor cortex.

While we identified a significant correlation between CD68 and pTDP-43 pathology in human ALS, one of the limitations of utilising post-mortem human tissue is the inability to assess changes temporally. We circumvented this issue by using a brain region less affected in disease or only affected in late-stage disease: the hippocampus, which shows minimal pathology in sALS, with only stage 4 cases developing pTDP-43 in the hippocampal dentate gyrus [50]. Interestingly, despite including within our ALS cohort stage 4 cases with pTDP-43 pathology in the hippocampal granule cell layer, we only identified microglial changes in the end-stage human ALS motor cortex, not hippocampus. Furthermore, the pattern of correlations between microglial functional marker expression in the ALS hippocampus mirrored that of the control brain. While previous studies investigating microglial CD68 expression in the ALS hippocampus did not compare to control cases, it has been noted that hippocampal CD68 expression did not correlate with pTDP-43 load, dementia state, or impaired executive function [41]. These data not only suggest that microglia in the ALS hippocampus share a phenotype with the control brain rather than the ALS motor cortex, but also that pTDP-43 is the primary pathology that subsequently drives microglial functional changes.

Based on the data from the end-stage human sALS tissue, we hypothesised that pTDP-43 aggregation is the primary pathology that triggers other neuropathological changes and microglial reactions. However, we have already identified that the major limitation of post-mortem human tissue work is the inability to study temporal changes in disease. To test our temporal hypothesis, we utilised the ‘rNLS’ TDP-43-driven model of ALS [51]. In this model, the removal of dietary DOX results in the neuronal expression of cytoplasmic human TDP-43 [51]. We observed this in our cohort, where pTDP-43 immunoreactivity was significantly increased in rNLS mice relative to controls from disease onset (2 WOD) and was maintained through to the late disease stage. We subsequently investigated microglial CD68 and L-ferritin changes relative to this pTDP-43 accumulation. Microglia increased their expression of CD68 at 4 WOD and L-ferritin at 6 WOD, supporting our temporal hypothesis that neuronal pTDP-43 pathology drives microglial changes. Further, this recapitulates the microglial phenotype shift hypothesised to occur in the human ALS brain, where a pool of phagocytic and neuroprotective CD68^high^ microglial shift to a dysfunctional L-ferritin^high^ state late in disease. It remains unclear what is driving the L-ferritin expression by microglia in the rNLS mice, and it is unclear whether iron dyshomeostasis occurs following TDP-43 accumulation. However, given the parallels between microglial changes in the rNLS mice and human ALS brain, pTDP-43 aggregation drives microglial changes in ALS either directly through microglial phagocytosis of pTDP-43 or indirectly through microglial responses to dysfunctional neurons.

Overall, we have demonstrated that microglia increase CD68 and L-ferritin expression following pTDP-43 aggregation. But the question remains: are reactive microglia neurotoxic or neuroprotective in the human ALS brain? The CD68^high^ microglial phenotype identified in the end-stage human motor cortex mirrors that of the neuroprotective microglia in the rNLS TDP-43-driven mouse model of ALS [18]. During the disease-recovery phase of late disease rNLS mice, there is an increase in microglial density, a shift to a more amoeboid morphology, and an increase in phagocytic *Cd68* gene expression. The improvement in motor symptoms in the rNLS mice during recovery following the microglial human TDP-43 uptake suggests that this phagocytic microglial activity was neuroprotective rather than neurotoxic [18]. Furthermore, microglial phagocytosis of damaged spinal motor neurons, including mislocalised intracellular TDP-43, has been demonstrated to be neuroprotective; the depletion of microglia enhanced neurodegeneration [68]. Finally, impairment of microglial phagocytosis through TREM2 deficiency enhances hTDP-43-overexpression induced neuronal degeneration, motor symptoms, and mortality [67]. Therefore, we postulate that the increase in microglial CD68 expression, and inferred phagocytosis, associated with pTDP-43 aggregation in both the rNLS mouse model of ALS and the human ALS brain is indeed neuroprotective.

Our data from both the human sALS and mouse rNLS brains suggest that a pool of neuroprotective CD68^high^ microglia shift into a dysfunctional L-ferritin^high^ state late in disease. Increased expression of iron storage genes at late-stage disease have already been described previously in the rNLS model [11]. While the debate regarding microglial functioning in ALS has largely been between whether neuroprotective or neurotoxic states develop, microglial dysfunction must also be considered. In the microglial dysfunction hypothesis of Alzheimer’s disease, microglia become progressively overwhelmed due to their reactive states in the aging and diseased brain [71]; in ALS, this could include the phagocytosis of pTDP-43 and degenerating neurons. In Alzheimer’s, overwhelmed microglia exhibit ‘frustrated phagocytosis’ and dysfunction, characterised by impaired phagocytosis, a dystrophic morphology, impaired migration, altered signalling, impaired proteostasis, and the development of senescence [71,72]. L-ferritin expression is specific for dystrophic microglia and, as such, demonstrates their presence in the ALS brain. Indeed, microglia derived from sALS patient monocytes have also been shown to have impaired phagocytic capacities which correlated with the rate of disease progression [73]. Together, this supports the hypothesis that a pool of microglia become dysfunctional in late-stage ALS. Therefore, therapeutics delivered early in disease should promote microglial phagocytic functions to maintain neuroprotection and reduce dysfunction, while those delivered in later stages should mitigate microglial dysfunction to maintain trophic support.

Despite evidence of microglia showing traits of neuroprotective and dysfunctional phenotypes, it remains possible that microglia acquire a neurotoxic phenotype in response to pTDP-43; a neurotoxic microglial phenotype has been induced *in vitro* in response to both native and mutant TDP-43 species [9]. Recent transcriptomic analyses of the human ALS spinal cord demonstrate that certain microglial functional states negatively correlated with disease duration, which was suggested to be evidence of microglial neurotoxicity in ALS [10]. However, the microglial state that negatively correlated with disease progression showed high expression of *CD68* and *FTL* (L-ferritin) [10]. While this could be a neurotoxic microglial state, we posit that, like in Alzheimer’s disease, this L-ferritin^high^ population is dysfunctional.

It remains unclear whether disease-associated changes in microglial function, and potential neuroprotective, dysfunctional, or neurotoxic effects, are consistent across ALS genotypes [42,44]. While pTDP-43 pathology occurs in 97% of ALS cases, mutations in *SOD1, C9orf72*, or *FUS* genes cause unique neuropathology that may drive distinct microglial responses. However, previous studies have shown similarities between microglial responses in *SOD1, C9orf72*, and *FUS*-linked ALS cases when compared with sALS [18,25]. Whether a microglial functional shift from CD68^high^ to L-ferritin^high^ is a consistent feature across ALS genotypes will determine the breath of utility of microglial-targeting therapies developed in TDP-43 models.

The evidence presented in this study demonstrates that microglia up-regulate proteins implicated in phagocytosis early in disease and dysfunction late in disease in ALS. We demonstrate that these microglial changes occur in response to pTDP-43 pathology and likely reflect transition from a neuroprotective role to a dysfunctional role for microglia in ALS pathogenesis.

## List of abbreviations

ALS: Amyotrophic lateral sclerosis
DOX: Doxycycline
GFAP: Glial fibrillary acidic protein
HLA-DR: Human leukocyte antigen, DR isotype
Iba1: Ionised calcium-binding adaptor protein 1
L-ferritin: Light chain ferritin
MOI: Marker of interest
pTDP-43: Phosphorylated TAR DNA-binding protein of 43 kDa
rNLS: Bigenic *NFFH*-tTA/*tetO*-hTDP-43ΔNLS mice
ROI: Region of interest
SOD1: Superoxide dismutase 1
TDP-43: TAR DNA-binding protein of 43 kDa
WOD: Weeks off doxycyclineDeclarations

## Ethics approval and consent to participate

All human brain tissue was donated to the Neurological Foundation Human Brain Bank following donor and donor family consent,and its use in this study was approved by the University of Auckland Human Participants Ethics committee (protocol number 011654). Animal ethics approval was obtained from The University of Queensland (#QBI/131/18), and experiments were conducted in accordance with the Australian code of practice for the care and use of animals for scientific purposes.

## Consent for publication

Not applicable.

## Availability of data and materials

The datasets used and/or analysed during the current study available from the corresponding author on reasonable request.

## Authors’ contributions

MEVS, AKW, and ELS conceptualized the study. RLMF and MAC collected and processed the human tissue. CT carried out pathological assessment of all cases. MEVS, MM, HCM, and MCC carried out the experimental procedures. MEVS and ELS developed the custom image analysis pipelines. MEVS and MM carried out quantification. Data analysis and interpretation were carried out by MEVS, MM, and ELS. Manuscript was prepared by MEVS and ELS. All authors read and approved of the final manuscript.

## Acknowledgements

This publication is dedicated to the incredible patients and families who contribute to our research. We would like to acknowledge Marika Eszes at the Centre for Brain Research (University of Auckland, New Zealand) for her technical role in the Neurological Foundation of New Zealand Human Brain Bank and the imaging team at the Biomedical Imaging Research Unit (University of Auckland, New Zealand). We would also like to thank Lidia Madrid San Martin, Wei Luan, Heledd Brown-Wright, and staff of the Queensland Brain Institute animal facility for technical and animal husbandry assistance.

## References

1. Mandrioli J, Faglioni P, Nichelli P, Sola P (2006) Amyotrophic lateral sclerosis: Prognostic indicators of survival. Amyotrophic Lateral Sclerosis 4: 217–226.

2. Scotter EL, Chen H-J, Shaw CE (2015) TDP-43 Proteinopathy and ALS: Insights into Disease Mechanisms and Therapeutic Targets. Neurotherapeutics 2: 352–363.

3. Al-Chalabi A, Calvo A, Chio A, Colville S, Ellis CM, et al. (2014) Analysis of amyotrophic lateral sclerosis as a multistep process: a population-based modelling study. Lancet Neurol 11: 1108–1113.

4. Neumann M, Sampathu DM, Kwong LK, Truax AC, Micsenyi MC, et al. (2006) Ubiquitinated TDP-43 in frontotemporal lobar degeneration and amyotrophic lateral sclerosis. Science 5796: 130–3.

5. Kettenmann H, Hanisch U-K, Noda M, Verkhratsky A (2011) Physiology of microglia. Physiol Rev 2: 461–553.

6. Liao B, Zhao W, Beers DR, Henkel JS, Appel SH (2012) Transformation from a neuroprotective to a neurotoxic microglial phenotype in a mouse model of ALS. Exp Neurol 1: 147–52.

7. Perry VH, Nicoll JAR, Holmes C (2010) Microglia in neurodegenerative disease. Nat Rev Neurol 4: 193–201.

8. Liao B, Zhao W, Beers DR, Henkel JS, Appel SH (2012) Transformation from a neuroprotective to a neurotoxic microglial phenotype in a mouse model of ALS. Exp Neurol 1: 147–52.

9. Zhao W, Beers DR, Bell S, Wang J, Wen S, et al. (2015) TDP-43 activates microglia through NF-κB and NLRP3 inflammasome. Exp Neurol: 24–35.

10. Humphrey J, Venkatesh S, Hasan R, Herb JT, de Paiva Lopes K, et al. (2023) Integrative transcriptomic analysis of the amyotrophic lateral sclerosis spinal cord implicates glial activation and suggests new risk genes. Nat Neurosci 1: 150–162.

11. Hunter M, Spiller KJ, Dominique MA, Xu H, Hunter FW, et al. (2021) Microglial transcriptome analysis in the rNLS8 mouse model of TDP-43 proteinopathy reveals discrete expression profiles associated with neurodegenerative progression and recovery. Acta Neuropathol Commun 1: 1–19.

12. Graber DJ, Hickey WF, Harris BT (2010) Progressive changes in microglia and macrophages in spinal cord and peripheral nerve in the transgenic rat model of amyotrophic lateral sclerosis. J Neuroinflammation: 1–12.

13. Stamenković S, Dučić T, Stamenković V, Kranz A, Andjus PR (2017) Imaging of glial cell morphology, SOD1 distribution and elemental composition in the brainstem and hippocampus of the ALS hSOD1G93A rat. Neuroscience: 37–55.

14. Sanagi T, Yuasa S, Nakamura Y, Suzuki E, Aoki M, et al. (2010) Appearance of phagocytic microglia adjacent to motoneurons in spinal cord tissue from a presymptomatic transgenic rat model of amyotrophic lateral sclerosis. J Neurosci Res 12: 2736–2746.

15. Gerber YN, Sabourin JC, Rabano M, Vivanco M d. M, Perrin FE (2012) Early functional deficit and microglial disturbances in a mouse model of amyotrophic lateral sclerosis. PLoS One 4.

16. Ohgomori T, Yamada J, Takeuchi H, Kadomatsu K, Jinno S (2016) Comparative morphometric analysis of microglia in the spinal cord of SOD1G93A transgenic mouse model of amyotrophic lateral sclerosis. European Journal of Neuroscience 10: 1340–1351.

17. Schludi MH, Becker L, Garrett L, Gendron TF, Zhou Q, et al. (2017) Spinal poly-GA inclusions in a C9orf72 mouse model trigger motor deficits and inflammation without neuron loss. Acta Neuropathol 2: 241–254.

18. Spiller KJ, Restrepo CR, Khan T, Dominique MA, Fang TC, et al. (2018) Microglia-mediated recovery from ALS-relevant motor neuron degeneration in a mouse model of TDP-43 proteinopathy. Nat Neurosci 3: 329–340.

19. Jara JH, Genç B, Stanford MJ, Pytel P, Roos RP, et al. (2017) Evidence for an early innate immune response in the motor cortex of ALS. J Neuroinflammation 1: 1–20.

20. Chiu IM, Morimoto ETA, Goodarzi H, Liao JT, O’Keeffe S, et al. (2013) A neurodegeneration-specific gene-expression signature of acutely isolated microglia from an amyotrophic lateral sclerosis mouse model. Cell Rep 2: 385–401.

21. Funikov SY, Rezvykh AP, Mazin P v., Morozov A v., Maltsev A v., et al. (2018) FUS(1-359) transgenic mice as a model of ALS: pathophysiological and molecular aspects of the proteinopathy. Neurogenetics 3: 189–204.

22. Beers DR, Zhao W, Liao B, Kano O, Wang J, et al. (2011) Neuroinflammation modulates distinct regional and temporal clinical responses in ALS mice. Brain Behav Immun 5: 1025–1035.

23. Mitchell JC, McGoldrick P, Vance C, Hortobagyi T, Sreedharan J, et al. (2013) Overexpression of human wild-type FUS causes progressive motor neuron degeneration in an age- and dose-dependent fashion. Acta Neuropathol 2: 273–288.

24. Kwan JY, Jeong SY, van Gelderen P, Deng HX, Quezado MM, et al. (2012) Iron accumulation in deep cortical layers accounts for MRI signal abnormalities in ALS: Correlating 7 tesla MRI and pathology. PLoS One 4.

25. Nolan M, Scott C, Gamarallage MP, Lunn D, Carpenter K, et al. (2020) Quantitative patterns of motor cortex proteinopathy across ALS genotypes. Acta Neuropathol Commun 1: 98.

26. Liu W, Venugopal S, Majid S, Ahn IS, Diamante G, et al. (2020) Single-cell RNA-seq analysis of the brainstem of mutant SOD1 mice reveals perturbed cell types and pathways of amyotrophic lateral sclerosis. Neurobiol Dis February: 104877.

27. Dachet F, Liu J, Ravits J, Song F (2019) Predicting disease specific spinal motor neurons and glia in sporadic ALS. Neurobiol Dis April: 104523.

28. Dols-Icardo O, Montal V, Sirisi S, López-Pernas G, Cervera-Carles L, et al. (2020) Motor cortex transcriptome reveals microglial key events in amyotrophic lateral sclerosis. Neurology - Neuroimmunology Neuroinflammation 5: e829.

29. Lopes KO, Sparks DL, Streit WJ (2008) Microglial dystrophy in the aged and Alzheimer’s disease brain is associated with ferritin immunoreactivity. Glia 10: 1048–1060.

30. Swanson MEV, Murray HC, Ryan B, Faull RLM, Dragunow M, et al. (2020) Quantitative immunohistochemical analysis of myeloid cell marker expression in human cortex captures microglia heterogeneity with anatomical context. Sci Rep 1: 11693.

31. McGeer PL, Itagaki S, McGeer EG (1988) Expression of the histocompatibility glycoprotein HLA-DR in neurological disease. Acta Neuropathol 6: 550–557.

32. Martinez FO, Gordon S (2014) The M1 and M2 paradigm of macrophage activation: time for reassessment. F1000Prime Rep March: 13.

33. Melief J, Koning N, Schuurman KG, van de Garde MDB, Smolders J, et al. (2012) Phenotyping primary human microglia: Tight regulation of LPS responsiveness. Glia 10: 1506–1517.

34. Walker DG, Lue L-FF (2015) Immune phenotypes of microglia in human neurodegenerative disease: challenges to detecting microglial polarization in human brains. Alzheimers Res Ther 1: 56.

35. Becker-Herman S, Arie G, Medvedovsky H, Kerem A, Shachar I (2005) CD74 is a member of the regulated intramembrane proteolysis-processed protein family. Mol Biol Cell 11: 5061–9.

36. Zotova E, Bharambe V, Cheaveau M, Morgan W, Holmes C, et al. (2013) Inflammatory components in human Alzheimer’s disease and after active amyloid-β42 immunization. Brain 9: 2677–2696.

37. Holness CL, Simmons DL (1993) Molecular cloning of CD68, a human macrophage marker related to lysosomal glycoproteins. Blood 6: 1607–1613.

38. Ohsawa K, Imai Y, Kanazawa H, Sasaki Y, Kohsaka S (2000) Involvement of Iba1 in membrane ruffling and phagocytosis of macrophages/microglia. J Cell Sci: 3073–84.

39. Imai Y, Ibata I, Ito D, Ohsawa K, Kohsaka S (1996) A novel gene iba1 in the major histocompatibility complex class III region encoding an EF hand protein expressed in a monocytic lineage. Biochem Biophys Res Commun 3: 855–862.

40. Ito D, Imai Y, Ohsawa K, Nakajima K, Fukuuchi Y, et al. (1998) Microglia-specific localisation of a novel calcium binding protein, Iba1. Molecular Brain Research 1: 1–9.

41. Brettschneider J, Libon DJ, Toledo JB, Xie SX, McCluskey L, et al. (2012) Microglial activation and TDP-43 pathology correlate with executive dysfunction in amyotrophic lateral sclerosis. Acta Neuropathol 3: 395–407.

42. Rifai OM, O’Shaughnessy J, Dando OR, Munro AF, Sewell MDE, et al. (2023) Distinct neuroinflammatory signatures exist across genetic and sporadic ALS cohorts. bioRxiv (Preprint).

43. Brettschneider J, Toledo JB, van Deerlin VM, Elman L, McCluskey L, et al. (2012) Microglial activation correlates with disease progression and upper motor neuron clinical symptoms in amyotrophic lateral sclerosis. PLoS One 6: 13–15.

44. Rifai OM, Longden J, O’Shaughnessy J, Sewell MDE, Pate J, et al. (2022) Random forest modelling demonstrates microglial and protein misfolding features to be key phenotypic markers in C9orf72-ALS. Journal of Pathology 4: 366–381.

45. Paolicelli RC, Sierra A, Stevens B, Tremblay ME, Aguzzi A, et al. (2022) Microglia states and nomenclature: A field at its crossroads. Neuron 21: 3458–3483.

46. Zhou Q, Mareljic N, Michaelsen M, Parhizkar S, Heindl S, et al. (2020) Active poly-GA vaccination prevents microglia activation and motor deficits in a C9orf72 mouse model. EMBO Mol Med 2: 1–13.

47. Aronica E, Baas F, Iyer A, ten Asbroek ALMA, Morello G, et al. (2015) Molecular classification of amyotrophic lateral sclerosis by unsupervised clustering of gene expression in motor cortex. Neurobiol Dis: 359–376.

48. Noristani HN, Sabourin JC, Gerber YN, Teigell M, Sommacal A, et al. (2015) Brca1 is expressed in human microglia and is dysregulated in human and animal model of ALS. Mol Neurodegener 1: 1–13.

49. Keren-Shaul H, Spinrad A, Weiner A, Matcovitch-Natan O, Dvir-Szternfeld R, et al. (2017) A Unique Microglia Type Associated with Restricting Development of Alzheimer’s Disease. Cell 7: 1276–1290.e17.

50. Brettschneider J, del Tredici K, Toledo JB, Robinson JL, Irwin DJ, et al. (2013) Stages of pTDP-43 pathology in amyotrophic lateral sclerosis. Ann Neurol 1: 20–38.

51. Walker AK, Spiller KJ, Ge G, Zheng A, Xu Y, et al. (2015) Functional recovery in new mouse models of ALS/FTLD after clearance of pathological cytoplasmic TDP-43. Acta Neuropathol 5: 643–660.

52. Waldvogel HJ, Curtis MA, Baer K, Rees MI, Faull RLM (2007) Immunohistochemical staining of post-mortem adult human brain sections. Nat Protoc 6: 2719–2732.

53. Wright AL, della Gatta PA, Le S, Berning BA, Mehta P, et al. (2021) Riluzole does not ameliorate disease caused by cytoplasmic TDP-43 in a mouse model of amyotrophic lateral sclerosis. European Journal of Neuroscience 6: 6237–6255.

54. Murray HC, Dieriks BV, Swanson ME v, Anekal PV, Turner C, et al. (2020) The unfolded protein response is activated in the olfactory system in Alzheimer’s disease. Acta Neuropathol Commun 1: 109.

55. Dieriks BV, Dean JM, Aronica E, Waldvogel HJ, Faull RLM, et al. (2018) Differential fatty acid-binding protein expression in persistent radial glia in the human and sheep subventricular zone. Dev Neurosci 2: 145–161.

56. Murray HC, Johnson K, Sedlock A, Highet B, Dieriks BV, et al. (2022) Lamina-specific immunohistochemical signatures in the olfactory bulb of healthy, Alzheimer’s and Parkinson’s disease patients. Commun Biol 1: 1–12.

57. Swanson MEV, Scotter EL, Murray HC, Turner C, Faull RLMM, et al. (2020) Identification of a dysfunctional microglial population in human Alzheimer’s disease cortex using novel single-cell histology image analysis. Acta Neuropathol Commun 1: 1–16.

58. Woolf Z, Swanson MEV, Smyth LC, Mee EW, Schweder P, et al. (2021) Single-cell image analysis reveals a protective role for microglia in glioblastoma. Neurooncol Adv 1.

59. Krijthe JH (2015) T-Distributed Stochastic Neighbor Embedding using Barnes-Hut Implementation. https://github.com/jkrijthe/Rtsne

60. van der Maaten L (2015) Accelerating t-SNE using tree-based algorithms. Journal of Machine Learning Research: 3221–3245.

61. Kobak D, Berens P (2019) The art of using t-SNE for single-cell transcriptomics. Nat Commun 1.

62. Cao J, Spielmann M, Qiu X, Huang X, Ibrahim DM, et al. (2019) The single-cell transcriptional landscape of mammalian organogenesis. Nature 7745: 496–502.

63. Belkina AC, Ciccolella CO, Anno R, Halpert R, Spidlen J, et al. (2019) Automated optimized parameters for T-distributed stochastic neighbor embedding improve visualization and analysis of large datasets. Nat Commun 1: 1–12.

64. Beygelzimer A, Kakadet S, Langford J, Arya S, Mount D, et al. (2019) FNN: Fast Nearest Neighbor Search Algorithms and Applications. https://cran.r-project.org/web/packages/FNN/index.html

65. Csardi G, Nepusz T (2006) The igraph software package for complex network research. InterJournal: 1695.

66. Streit WJ, Walter SA, Pennell NA (1999) Reactive microgliosis. Prog Neurobiol 6: 563–581.

67. Xie M, Liu YU, Zhao S, Zhang L, Bosco DB, et al. (2022) TREM2 interacts with TDP-43 and mediates microglial neuroprotection against TDP-43-related neurodegeneration. Nat Neurosci 1: 26–38.

68. Svahn AJ, Don EK, Badrock AP, Cole NJ, Graeber MB, et al. (2018) Nucleo-cytoplasmic transport of TDP-43 studied in real time: impaired microglia function leads to axonal spreading of TDP-43 in degenerating motor neurons. Acta Neuropathol 3: 445–459.

69. Streit WJ, Braak H, Xue QS, Bechmann I (2009) Dystrophic (senescent) rather than activated microglial cells are associated with tau pathology and likely precede neurodegeneration in Alzheimer’s disease. Acta Neuropathol 4: 475–485.

70. Streit WJ, Xue QS (2009) Life and death of microglia. Journal of Neuroimmune Pharmacology 4: 371–379.

71. Streit WJ, Xue Q-S, Tischer J, Bechmann I (2014) Microglial pathology. Acta Neuropathol Commun 1: 142.

72. Mosher KI, Wyss-Coray T (2014) Microglial dysfunction in brain aging and Alzheimer’s disease. Biochem Pharmacol 4: 594–604.

73. Quek H, Cuní-López C, Stewart R, Colletti T, Notaro A, et al. (2022) ALS monocyte-derived microglia-like cells reveal cytoplasmic TDP-43 accumulation, DNA damage, and cell-specific impairment of phagocytosis associated with disease progression. J Neuroinflammation 1: 1–21.

