## Supplementary materials for "Microglial CD68 and L-ferritin upregulation in response to phosphorylated-TDP-43 pathology in the amyotrophic lateral sclerosis brain"

**Table S1:** Summary of microglial functional markers and changes identified in the CNS from human ALS patients and rodent models of ALS

| **Iba1** (Calcium-binding protein for membrane ruffling and phagocytosis expressed by all microglia [38–40]) | | | | | |
|---|---|---|---|---|---|
| **Species** | **Genotype/mutation** | **Region** | **Method** | **Microglial change** | **Reference** |
| Mouse | $hFUS^{1\text{-}359}$ | Cervical spinal cord | RNAseq | *Aif1* enriched in D120 mice relative to controls and in D60 mice relative to D120. | [21] |
| Rat | $hSOD1^{G93A}$ | Spinal cord | IHC | Iba1+ cells appeared in ventral horns of pre-clinical rats as focal aggregates, near but not juxtaposing NeuN-positive motor neurons. | [12] |
| Rat | $hSOD1^{G93A}$ | Hippocampus (dentate gyrus) | IHC | Progressive significant increase in the percentage of Iba1+ pixels in pre-symptomatic to symptomatic ALS rats, compared with age-matched controls.<br>This was not accompanied by any microglial morphological changes. | [13] |
| | | Facial nucleus | | Progressive significant increase in the percentage of Iba1+ pixels in pre-symptomatic to symptomatic ALS rats, compared with age-matched controls.<br>Microglial morphology was also changed; microglial surface area and volume were significantly reduced in pre-symptomatic and symptomatic mice relative to controls. | |
| Rat | $hSOD1^{H46R}$ | Lumbar spinal cord | IHC | Progressively increased Iba1 immunoreactivity in anterior horn of pre-symptomatic to early symptomatic transgenic rats, relative to controls.<br>Iba1+ microglia appeared around motor neurons. | [14] |
| Mouse | $hSOD1^{G93A}$ | Spinal cord | FACS | Microglia split into $Iba1^{low}$ and $Iba1^{high}$ populations in P60 transgenic mice; the percentage of $Iba1^{high}$ microglia was increased in transgenic mice relative to controls. | [15] |
| | | | IHC | Increased Iba1+ cell density in ventral horn of P60 mice, which exhibited more 'activated' morphology with larger somata and reduced processes. | |
| Mouse | $hSOD1^{G93A}$ | Lumbar spinal cord | IHC | Spatial distribution of microglia was regular at 9-weeks of age, but it became irregular in 15- and 21-week-old transgenic mice relative to age-matched controls.<br>Spatial changes were accompanied by morphological changes including but not limited to increased process number, and reduced node number and process length. | [16] |

| | | | | | |
|---|---|---|---|---|---|
| Mouse | *C9orf72* ((GA)$_{149}$-CFP) | Spinal cord | qPCR | Significant increase in *Aif1* at 1 and 6 months of age. | [17] |
| Mouse | rNLS-TDP-43 (*NEFH*-tTA/*tetO*-hTDP-43ΔNLS) | Spinal cord | IHC | Significant increase in Iba1+ cell density at mid- to late-stage disease (6 weeks off DOX) relative to age-matched controls. | [18] |
| Human | sALS | Lumbar spinal cord | IHC | Average microglial score (encompassing cell number and area coverage) was not consistently increased, relative to controls. | [18] |
| Human | *C9orf72* | Lumbar spinal cord | IHC | Average microglial score (encompassing cell number and area coverage) was not consistently increased, relative to controls. | [18] |
| Human | *SOD1* | Lumbar spinal cord | IHC | Average microglial score (encompassing cell number and area coverage) was significantly increased, relative to controls. | [18] |
| Human | sALS<br>*SOD1* (fALS) | Motor cortex | IHC | No difference in microglial cell density, but increased number of 'activated' microglia. | [19] |
| **CD68** (Lysosomal-associated protein upregulated in actively phagocytic cells [36,37]) | | | | | |
| **Species** | **Genotype/mutation** | **Region** | **Method** | **Microglial change** | **Reference** |
| Mouse | hSOD1$^{G93A}$ | Spinal cord | RNAseq | *Cd68* enriched in both D100 and D130 (end-stage disease) mice relative to age-matched controls. | [20] |
| Mouse | hSOD1$^{G93A}$ | Lumbar spinal cord | Microarray | *Cd68* enriched in D90 disease mice relative to age-matched controls. | [48] |
| Mouse | *C9orf72* ((GA)$_{149}$-CFP) | Thoracic spinal cord | Transcriptomic analysis | *Cd68* enriched in transgenic mice relative to age-matched controls. | [17] |
| Mouse | hFUS$^{1-359}$ | Cervical spinal cord | RNAseq | *Cd68* enriched in D120 mice relative to controls and in D60 mice relative to D120. | [21] |
| Mouse | hSOD1$^{G93A}$ | Cervical spinal cord | RT-qPCR | *Cd68* progressively increased from 16 weeks of age through until end-stage disease. | [22] |
| Mouse | hSOD1$^{G93A}$ | Cervical spinal cord | IHC | Immunoreactivity progressively increased from 11 weeks of age through until end-stage disease. | [22] |
| Mouse | hSOD1$^{G93A}$ | Lumbar spinal cord | RT-qPCR | *Cd68* progressively increased from 16 weeks of age through until end-stage disease. | [22] |
| Mouse | hSOD1$^{G93A}$ | Lumbar spinal cord | IHC | Immunoreactivity progressively increased from 11 weeks of age through until end-stage disease. | [22] |
| Mouse | hFUS | Spinal cord | IHC | Increased CD68 immunoreactivity with increased hFUS expression. | [23] |
| Mouse | *C9orf72* ((GA)$_{149}$-CFP) | Spinal cord | qPCR | Increased *Cd68* expression at 1 and 6 months of age. | [17] |

| | | | | | |
|---|---|---|---|---|---|
| Mouse | rNLS-TDP-43 (*NEFH*-tTA/*tetO*-hTDP-43ΔNLS) | Spinal cord | IHC | No change in CD68 immunoreactivity area coverage at any disease stage. | [18] |
| Rat | $hSOD1^{G93A}$ | Spinal cord | IHC | Increased CD68 immunoreactivity in ventral horn in symptomatic, but not pre-symptomatic animals. | [12] |
| Human | sALS | Motor cortex | IHC | CD68 immunoreactivity increased in deep cortical layers. | [24] |
| Human | *C9orf72* | Lumbar spinal cord | IHC | Increase in CD68+ cell density in cerebrospinal tract and anterior horn relative to control, but not significant. | [25] |
| | *SOD1* | | | Increase in CD68+ cell density in cerebrospinal tract and anterior horn relative to control, but not significant. | |
| | *FUS* | | | Increase in CD68+ cell density in cerebrospinal tract and anterior horn relative to control, but not significant. | |
| | sALS | | | Significant increase in CD68+ cell density in cerebrospinal tract and anterior horn relative to control.<br>CD68+ cell density did not correlate with pTDP-43 load. | |
| **L-ferritin (**Protein that sequesters free iron to reduce reactive oxygen species formation which is up-regulated by degenerating microglia [29,69]) | | | | | |
| **Species** | **Genotype/mutation** | **Region** | **Method** | **Microglial change** | **Reference** |
| Mouse | $hFUS^{1-359}$ | Cervical spinal cord | RNAseq | *Ftl1* and *ftl2* enriched in D60 mice relative to D120. | [21] |
| Human | sALS | Motor cortex | IHC | Ferritin immunoreactivity increased in areas of high iron by CD68+ microglia. | [24] |
| **HLA-DR** (Antigen presentation molecule considered the classical 'activation' marker [31–34]) | | | | | |
| **Species** | **Genotype/mutation** | **Region** | **Method** | **Microglial change** | **Reference** |
| Mouse | $hSOD1^{G93A}$ | Pontine brainstem | Drop sequencing | *H2-Eb1* enriched in D100 *SOD1* mice relative to age-matched controls | [26] |
| Mouse | rNLS-TDP-43 (*NEFH*-tTA/*tetO*-hTDP-43ΔNLS) | Spinal cord | scRNAseq | *H2-Eb1* progressively more enriched in early to late diseased mice relative to age-matched controls. | [18] |
| Rat | $hSOD1^{G93A}$ | Spinal cord | IHC | Increased HLA-DR in ventral grey matter after symptom onset, but not in pre-symptomatic rats. | [12] |
| Rat | $hSOD1^{G93A}$ | Lumbar spinal cord | IHC | Increased immunoreactivity on aggregated microglia in pre-symptomatic rats. | [14] |

| | | | | | |
|---|---|---|---|---|---|
| Human | sALS | Lumbar spinal cord | Laser capture microdissection and exon array technology | *HLA-DRA* enriched in 'M2' microglia/macrophages associated with lower motor neurons. | [27] |
| Human | sALS | Motor cortex | RNAseq | *HLA-DRA* enriched in ALS versus control. | [28] |
| **CD74** (HLA-DR invariant chain that supports HLA-DR formation and transportation to the cell surface [35]) | | | | | |
| **Species** | **Genotype/mutation** | **Region** | **Method** | **Microglial change** | **Reference** |
| Mouse | SOD1$^{G93A}$ | Spinal cord | RNAseq | *Cd74* enriched in D100 *SOD1* mice relative to age-matched control. | [20] |
| Mouse | rNLS-TDP-43 (*NEFH*-tTA/*tetO*-hTDP-43ΔNLS) | Spinal cord | scRNAseq | *Cd74 e*nriched in late disease relative to control | [18] |
| Mouse | SOD1$^{G93A}$ | Pontine brainstem | Drop sequencing | Depleted microglial expression of *cd74* in D100 *SOD1* mice relative to age-matched control. | [26] |
| Human | sALS | Lumbar spinal cord | Laser capture microdissection and exon array technology | *Cd74 e*nriched in 'M2' microglia/macrophages associated with lower motor neurons. | [27] |
| Human | sALS | Motor cortex | RNAseq | *Cd74 e*nriched in ALS versus control. | [28] |

Abbreviations: FACS, fluorescence-activated cell sorting; IHC, immunohistochemistry; qPCR, quantitative polymerase chain reaction; RT-qPCR, quantitative reverse transcription PCR; RNAseq, RNA sequencing; sALS, sporadic amyotrophic lateral sclerosis; scRNAseq, single-cell RNA sequencing; SOD1, superoxide dismutase 1; TDP-43, TAR DNA-binding protein of 43 kDa.

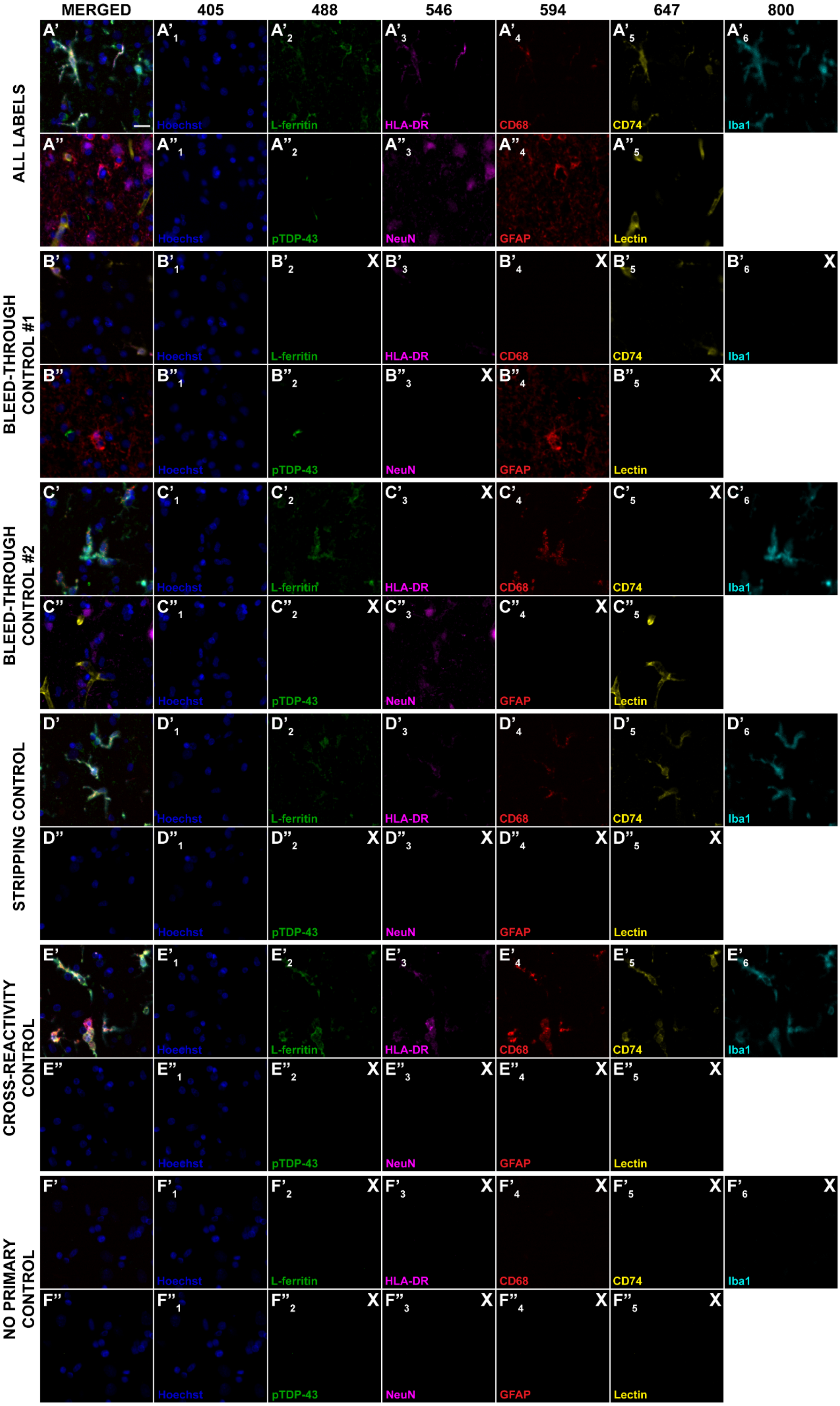

MERGED
405
488
546
594
647
800
ALL LABELS
BLEED-THROUGH CONTROL #1
BLEED-THROUGH CONTROL #2
STRIPPING CONTROL
CROSS-REACTIVITY CONTROL
NO PRIMARY CONTROL
Hoechst
L-ferritin
HLA-DR
CD68
CD74
Iba1
pTDP-43
NeuN
GFAP
Lectin

**Figure S1: Two-round multiplexed immunohistochemistry shows no bleed-through or cross-reactivity within and between rounds**

Two rounds of immunohistochemical labelling were used to visualise microglial, anatomical, and pathological markers in the motor cortex and hippocampus from human ALS and neurologically normal control cases. To investigate bleed-through and cross-reactivity within and between immunohistochemical labelling rounds, different labelling conditions were tested: all labels with all primary and secondary antibodies in both rounds (**A**), all primary antibodies with alternating secondary antibodies omitted in both rounds (indicated by an X) to test channel bleed through (**B and C**), a stripping control where all primary and secondary antibodies were applied for round 1, but no antibodies were applied after stripping (as indicated by an X) (**D**), a cross-reactivity control where all primary and secondary antibodies were applied for round 1, and only secondary antibodies were applied for round 2 (**E**), and a no primary control where no primary or secondary antibodies were applied for either labelling round (**F**). Round 1 (denoted by **'**) was comprised of microglial markers, including L-ferritin (green; 2), HLA-DR (magenta; 3), CD68 (red; 4), CD74 (yellow; 5), and Iba1 (cyan;6), with a Hoechst nuclear counterstain (blue; 1). Round 2 (denoted by **"**) was comprised of pathological and anatomical markers, including pTDP-43 (green; 2), NeuN (magenta; 3), GFAP (red; 4), and lectin (yellow; 5) with a Hoechst nuclear counterstain (blue; 1). Images from rounds 1 and 2 were aligned using the Hoechst nuclear counterstain. Scale bar = 20 µm.

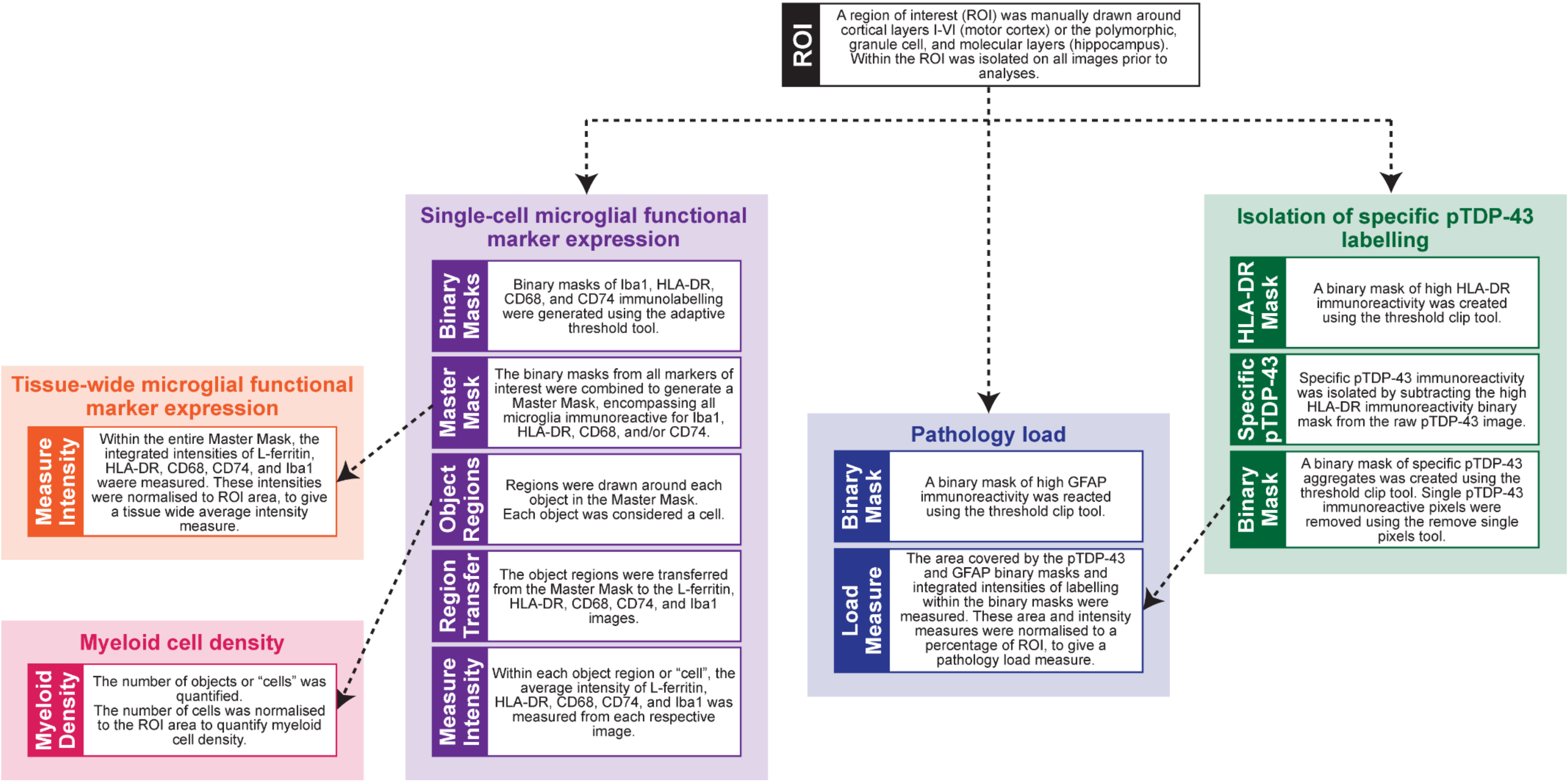


**Figure S2: Flow diagram summarising Metamorph images analysis pipelines**

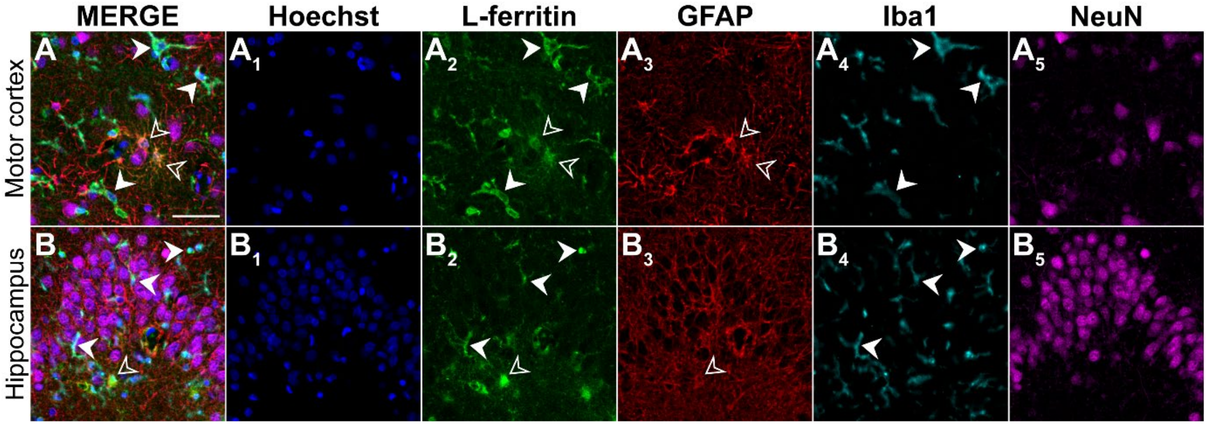


**Figure S3: Astrocytic expression of L-ferritin in the human brain**

Post-mortem human motor cortex (**A**) and hippocampus (**B**) tissue was immunolabelled with a Hoechst nuclear counterstain (blue; 1), L-ferritin (green; 2), GFAP (red; 3), Iba1 (cyan; 4), and NeuN (magenta; 5). L-ferritin-positive GFAP-positive astrocytes are indicated with filled-in arrows and L-ferritin-positive Iba1-positive Iba1 microglia are indicated with hollow arrows. Scale bar = 20 µm.

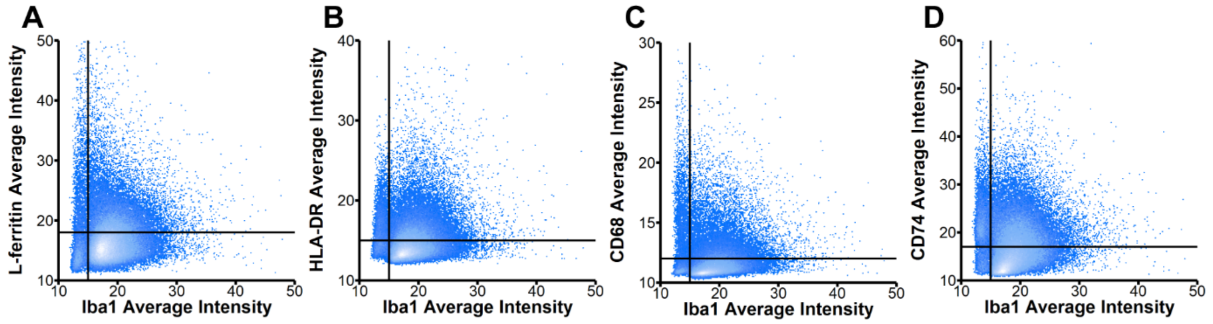


**Figure S4: Iba1-marker of interest scatter plots to determine high-low thresholds**

Single-cell microglial average intensities of L-ferritin, HLA-DR, CD68, CD74, and Iba1 were quantified in the motor cortex and hippocampus from control and ALS cases using novel Metamorph image analysis pipelines. When all microglia from all control and ALS cases were plotted as a distribution curve for average intensity of each marker, only Iba1 had distinct low and high-expressing populations (Figure 3N). To determine the threshold for low- and high-expressing populations for the other microglial functional markers, each microglia from all control and ALS cases was plotted on an *XY* scatter plot for Iba1 average intensity against L-ferritin (A), HLA-DR (B), CD68 (C), and CD74 (D). The threshold for Iba1 is indicated by a vertical line on each plot, while the threshold for each function marker is indicated by a horizontal line on each plot.

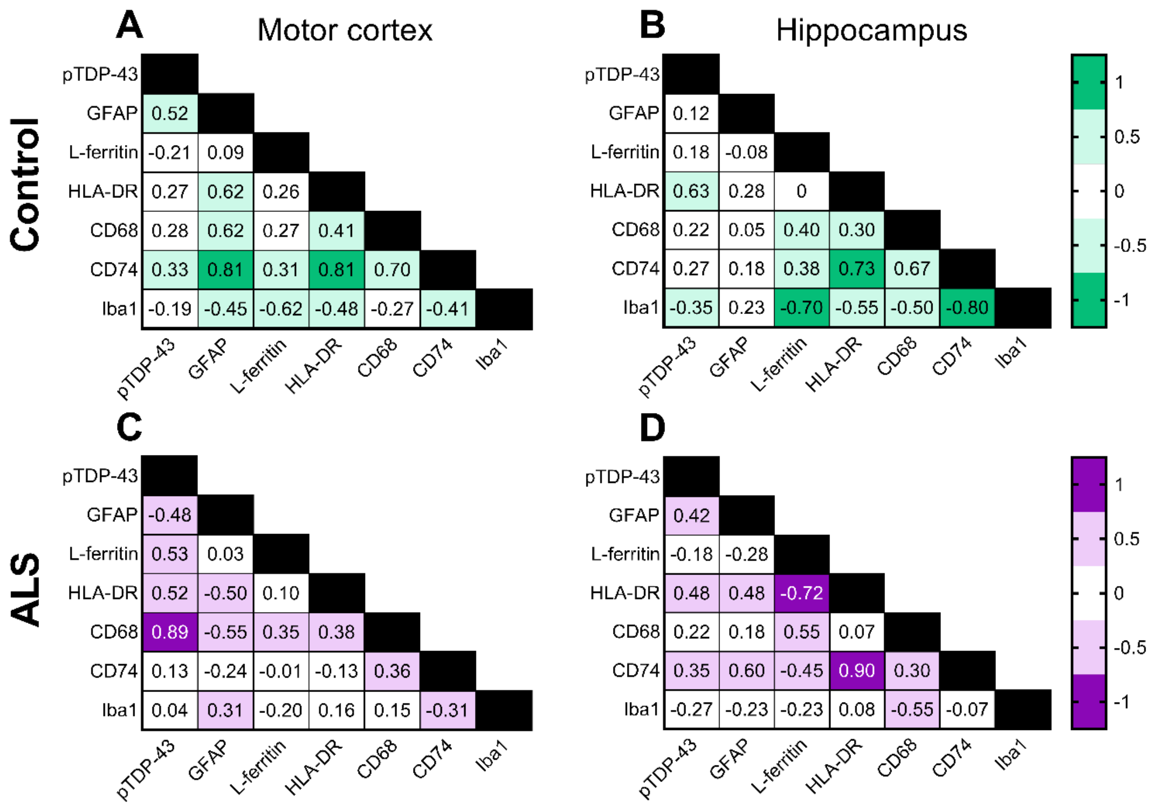


**Figure S5: Microglial CD68 average intensity correlates with pTDP-43 load in the ALS motor cortex**

Measures of microglial functional changes and neuropathology load were correlated in the control (**A-B**) and ALS (**C-D**) motor cortex and hippocampus. Measures of pTDP-43 integrated intensity and GFAP area coverage, and microglial average intensities of L-ferritin, HLA-DR, CD68, CD74, and Iba1 microglia were sequentially correlated with one another using Spearman correlations. The resulting r value from each correlation is presented in the correlation matrix and colour coded relative to strength. All r values greater than 0.7 were statistically significant and considered very strong correlations. Control n = 10 and ALS n = 9-10, where statistical significance was set at $p \leq 0.05$.

| **Table S2: Correlations between motor cortex pathology and microglial measures with age and post-mortem delay** | | | | |
|---|---|---|---|---|
| **Age** | | | | |
| | **ALS** | | **Control** | |
| | **r** | **P value** | **r** | **P value** |
| **pTDP-43** | -0.5636 | 0.0963 | -0.4268 | 0.2192 |
| **GFAP** | 0.5273 | 0.1231 | 0.06098 | 0.8714 |
| **L-ferritin$^{high}$** | -0.02432 | 0.9528 | 0.1951 | 0.5873 |
| **HLA-DR$^{high}$** | -0.2067 | 0.5645 | 0.3781 | 0.2799 |
| **CD68$^{high}$** | -0.4499 | 0.1935 | 0.3171 | 0.3703 |
| **CD74$^{high}$** | -0.2675 | 0.4529 | 0.2256 | 0.5297 |
| **Iba1$^{high}$** | -0.1459 | 0.6881 | -0.3110 | 0.3785 |
| **Post-mortem delay** | | | | |
| | **ALS** | | **Control** | |
| | **r** | **P value** | **r** | **P value** |
| **pTDP-43** | 0.1945 | 0.5892 | 0.6970 | 0.0306 |
| **GFAP** | -0.4012 | 0.2506 | 0.2970 | 0.4069 |
| **L-ferritin$^{high}$** | 0.4207 | 0.2244 | 0.006061 | >0.9999 |
| **HLA-DR$^{high}$** | 0.09146 | 0.8008 | 0.3091 | 0.3869 |
| **CD68$^{high}$** | 0.2134 | 0.5498 | 0.4909 | 0.1548 |
| **CD74$^{high}$** | 0.4268 | 0.2178 | 0.1394 | 0.7072 |
| **Iba1$^{high}$** | -0.8476 | 0.0030 | -0.3333 | 0.3487 |

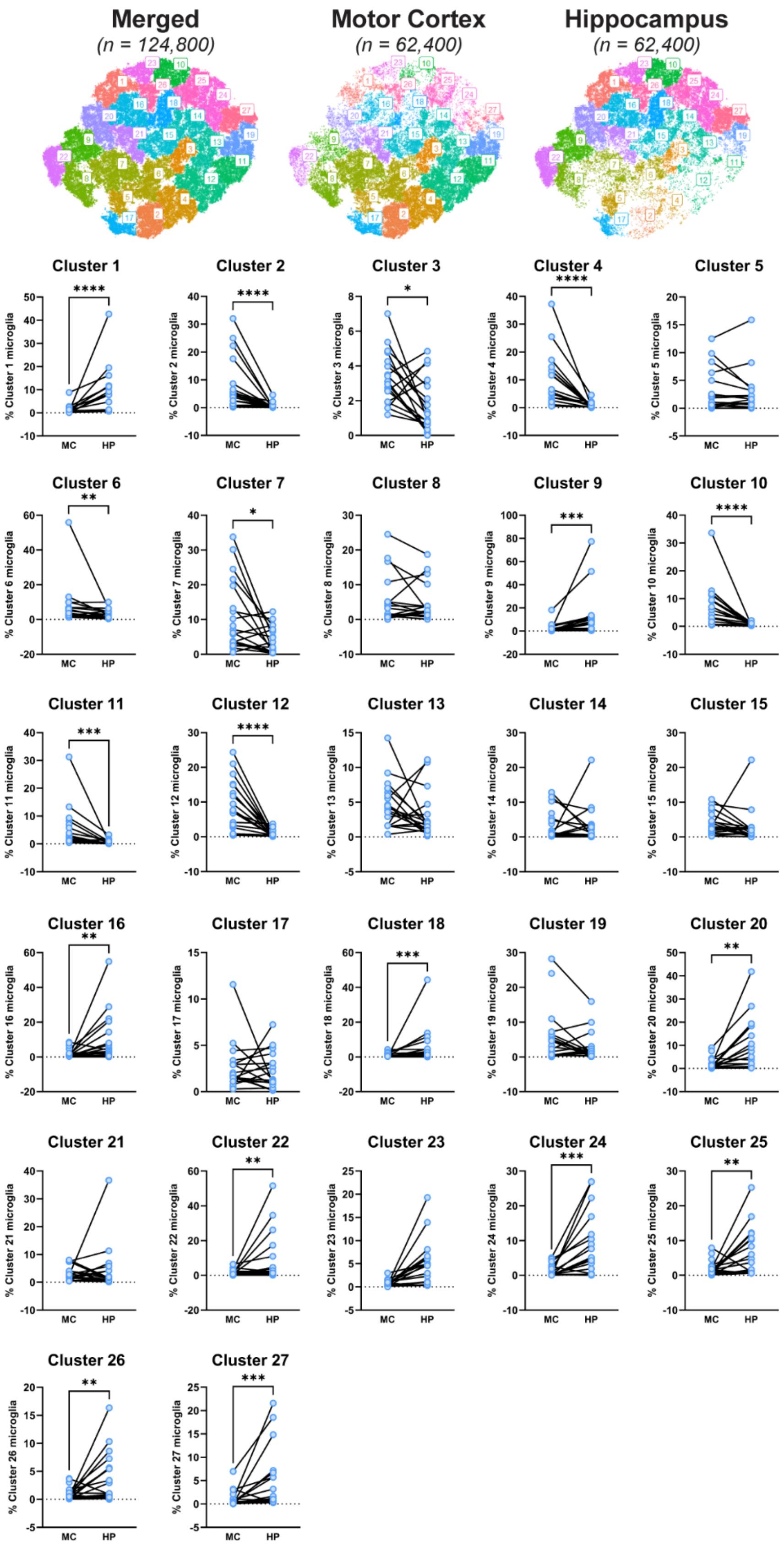


**Figure S6: Microglial clusters are differentially abundant in the motor cortex and hippocampus of the human brain**

Microglial single-cell average intensities of L-ferritin, HLA-DR, CD68, CD74, and Iba1 were used to identify and characterise microglial subpopulations that were changed in ALS relative to controls. A t-SNE plot was generated using random subset of 124,800 microglia (31,200 each from control motor cortex, ALS motor cortex, control hippocampus, and ALS hippocampus), and 27 clusters were identified using a k nearest neighbour and Louvain clustering approach. The percentage of motor cortex (MC) and hippocampus (HP) microglia in each cluster was determined for each case, irrespective of case group. Data presented with each case represented by a single point and the percentages of microglia in each cluster in the motor cortex and hippocampus are joined per case. The percentage of microglia in each cluster in each region was compared between motor cortex and hippocampus using Wilcoxon matched-pairs signed rank tests. Significance of differences between regions: ****$p \leq 0.0001$, *** $p \leq 0.001$, **$p \leq 0.01$, * $p \leq 0.05$.

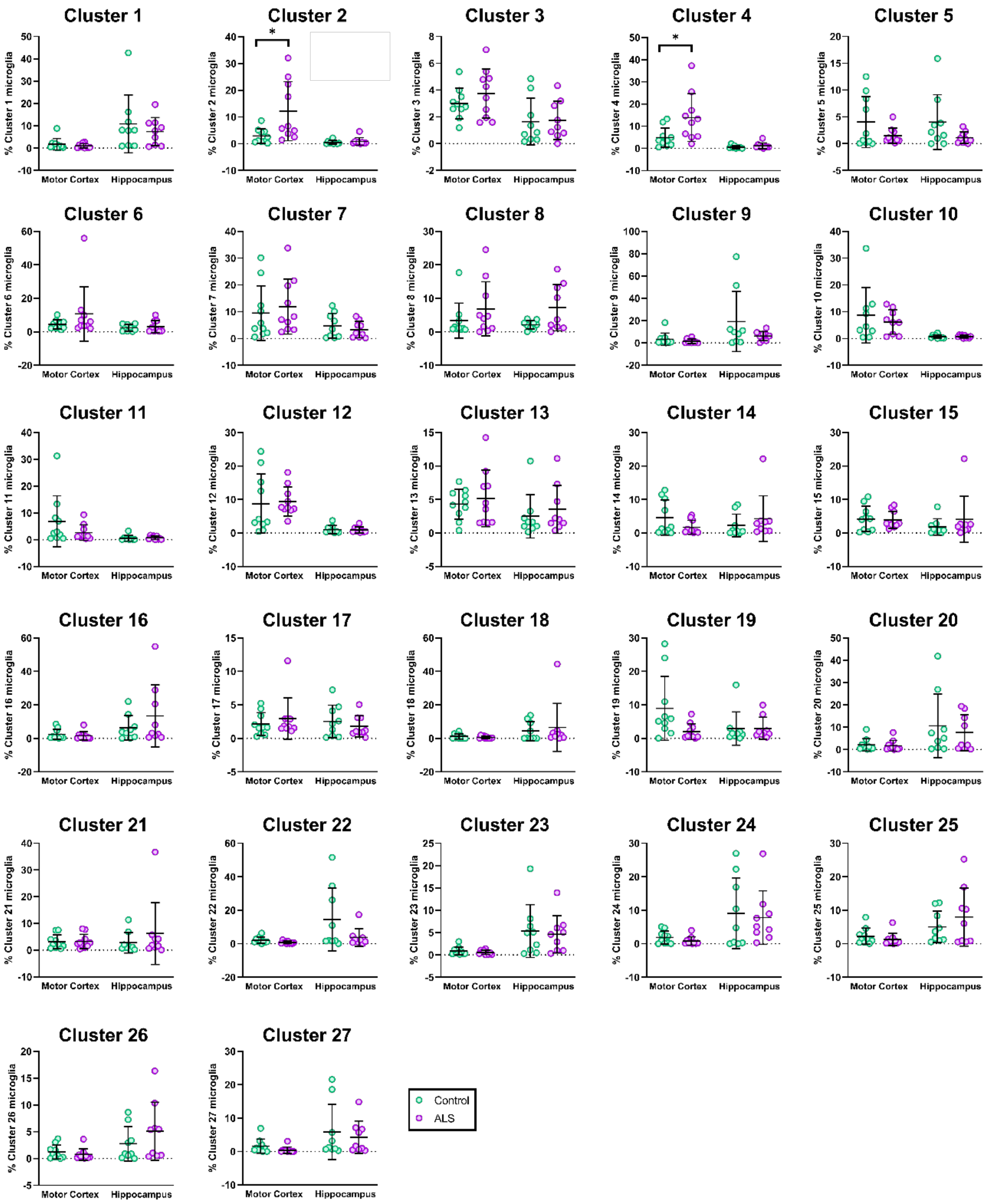


**Figure S7: Microglial clusters are differentially abundant in the human ALS brain.**

Microglial single-cell average intensities of L-ferritin, HLA-DR, CD68, CD74, and Iba1 were used to identify and characterise microglial subpopulations that were changed in ALS cases relative to controls. A t-SNE plot was generated using random subset of 124,800 microglia (31,200 each from control motor cortex, ALS motor cortex, control hippocampus, and ALS hippocampus), and 27 clusters were identified using a k nearest neighbour and Louvain clustering approach (**Fig 6A**). The percentage of total microglia in each of the 27 clusters were statistically compared between control and ALS motor cortex and hippocampus. Data presented as mean ± SD; control n = 10 and ALS n = 9-10. Cluster proportions were compared between case groups with multiple Mann-Whitney tests and multiple comparisons were controlled for using a False Discovery Rate of 0.01, as determined by the two-stage step-up method of Benjamini, Krieger, and Yekutieli. Significance of differences between case groups: *$p \leq 0.05$.

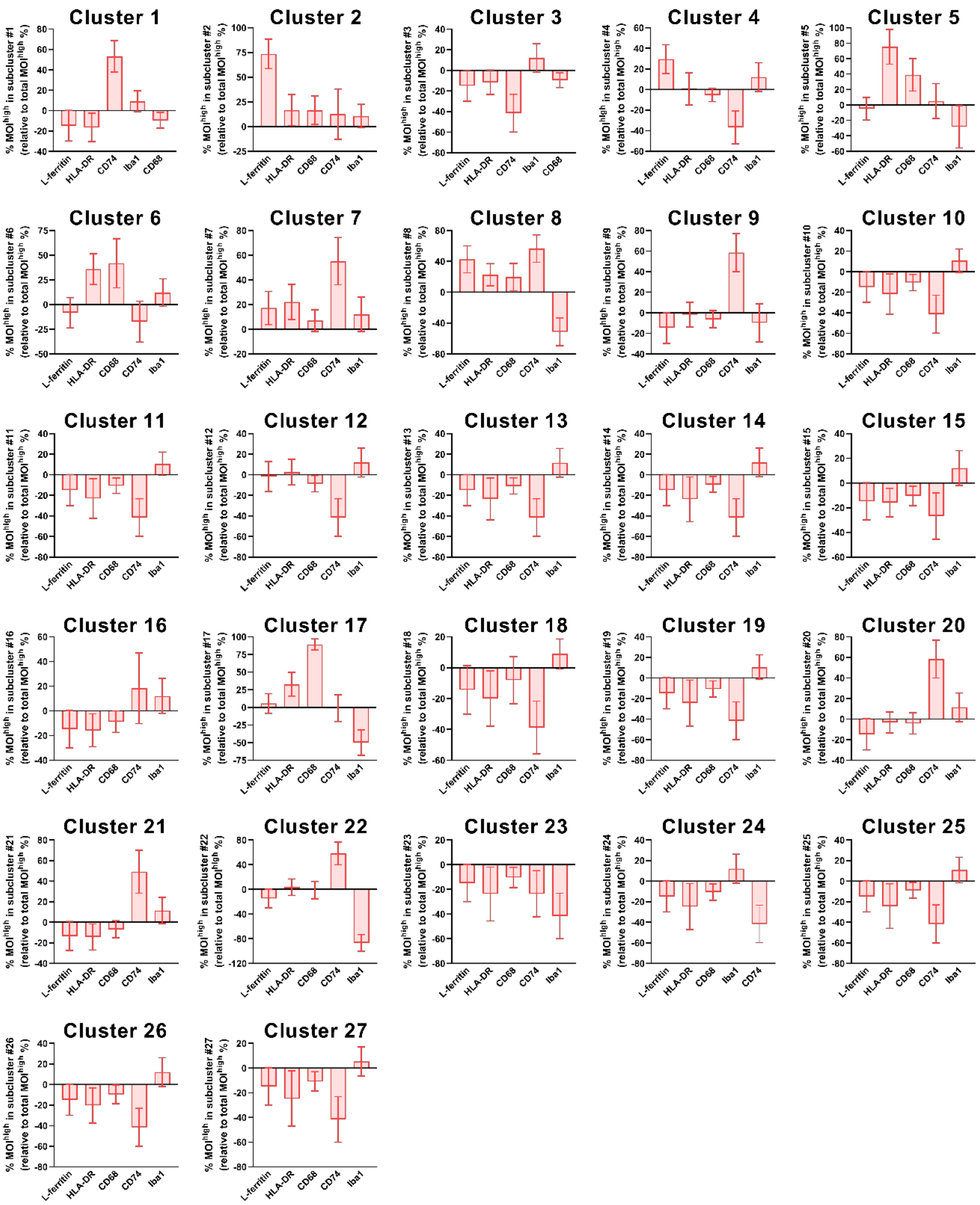


**Figure S8: Relative percentage of L-ferritin, HLA-DR, CD68, CD74, and Iba1 high-expressing microglia in each cluster**

Microglial single-cell average intensities of L-ferritin, HLA-DR, CD68, CD74, and Iba1 were used to identify and characterise microglial subpopulations identified in ALS cases relative to controls. The percentage high-expressing microglia were determined per cluster for each functional marker of interest. The percentage $MOI^{high}$ for each functional marker was normalised to the total percentage of $MOI^{high}$ in each case for cluster 1 to 27 in all cases. Data presented as mean ± SD; n = 20.
